# A single N6-methyladenosine site in lncRNA HOTAIR regulates its function in breast cancer cells

**DOI:** 10.1101/2020.06.08.140954

**Authors:** Allison M. Porman, Justin T. Roberts, Emily D. Duncan, Madeline L. Chrupcala, Ariel A. Levine, Michelle A. Kennedy, Michelle M. Williams, Jennifer K. Richer, Aaron M. Johnson

**Author notes:** Corresponding Author Aaron M. Johnson, PhD.

## Abstract

N6-methyladenosine (m6A) modification of RNA plays important roles in normal and cancer biology, but knowledge of its function on long noncoding RNAs (lncRNAs) remains limited. Here, we investigate whether m6A regulates the function of the human HOTAIR lncRNA, which contributes to multiple pro-tumor phenotypes in triple-negative breast cancer (TNBC) cells. We identify at least 8 individual m6A sites within HOTAIR, with a single site (A783) consistently methylated. Mutation of A783 impairs cellular proliferation and invasion in HOTAIR-overexpressing TNBC cells. m6A at A783 regulates HOTAIR’s ability to localize to chromatin and induce gene pathways that affect tumor progression. In contrast, A783U mutant HOTAIR demonstrates loss-of-function and antimorph behaviors by impairing gene expression changes induced by WT HOTAIR and, in some cases, inducing opposite changes in gene expression. HOTAIR interacts with nuclear m6A reader YTHDC1 and high HOTAIR is significantly associated with shorter overall patient survival, particularly in the context of high *YTHDC1*. At the molecular level, YTHDC1-HOTAIR interactions are required for chromatin localization and regulation of gene repression. Our work demonstrates how modification of one base in a lncRNA can elicit a distinct gene regulation mechanism and drive disease-associated phenotypic changes such as proliferation and invasion.

## Introduction

Long non-coding RNAs (lncRNAs) are becoming increasingly noted for their roles in transcriptional regulation(Long, Wang, Youmans, & Cech, 2017). Members of this class of noncoding RNAs are typically longer than 200 nucleotides, transcribed by RNA polymerase II, and processed similarly to mRNAs(Esteller, 2011). LncRNAs regulate transcription in a variety of ways; they can alter chromatin by directing histone-modifying enzymes to their target loci to induce changes in chromatin, or can regulate transcription directly by interacting with transcription factors and RNA polymerase II(Long et al., 2017). Importantly, lncRNAs are often key regulators of epigenetic changes that can drive cancer progression, often by aberrant overexpression(Schmitt & Chang, 2016).

The human lncRNA HOTAIR is a 2.2kb spliced and polyadenylated RNA transcribed from the HoxC locus. Originally identified as a developmental regulator acting in *trans* to repress expression of the HoxD locus (Rinn et al., 2007), aberrant high levels of HOTAIR are associated with poor survival and increased cancer metastasis in many different cancer types, including breast cancer(Balas & Johnson, 2018; Gupta et al., 2010). Exogenous overexpression of HOTAIR in the MDA-MB-231 TNBC cell line results in the repression of hundreds of genes(Gupta et al., 2010), and it promotes cell invasion, migration, proliferation, and self-renewal capacity in multiple breast cancer cell lines(Deng et al., 2017; Gupta et al., 2010; Meredith, Balas, Sindy, Haislop, & Johnson, 2016). HOTAIR function is particularly striking in MDA-MB-231 cells, given that this is already a highly invasive breast cancer cell line and its invasiveness is increased even further by HOTAIR overexpression(Gupta et al., 2010; Meredith et al., 2016). This is reflective of the prognostic impact of HOTAIR expression in TNBC patients where high HOTAIR expression correlates with poorer overall survival(Gupta et al., 2010; Yang et al., 2011). MDA-MB-231 cells express low levels of endogenous HOTAIR, offering an opportunity to study response to HOTAIR transgenic overexpression, which is proposed to mimic the high levels of HOTAIR observed in patients with aggressive TNBC(Gupta et al., 2010).

At its target loci, HOTAIR mediates the induction of H3K27 trimethylation (H3K27me3) by Polycomb Repressive Complex 2 (PRC2), resulting in heterochromatin formation and repression(Gupta et al., 2010; Tsai et al., 2010; Yansheng Wu et al., 2015). In cancer contexts, high levels of HOTAIR misdirect this mechanism to loci that are not typically repressed in the tissue of origin(Balas & Johnson, 2018; Gupta et al., 2010; Hajjari & Salavaty, 2015). Despite these previous findings, a recent study demonstrated that HOTAIR can repress genes even in the absence of PRC2, suggesting that initial repression or transcriptional interference may occur upstream of H3K27me3 by PRC2(Portoso et al., 2017) (Figure 1A).

**Figure 1.**
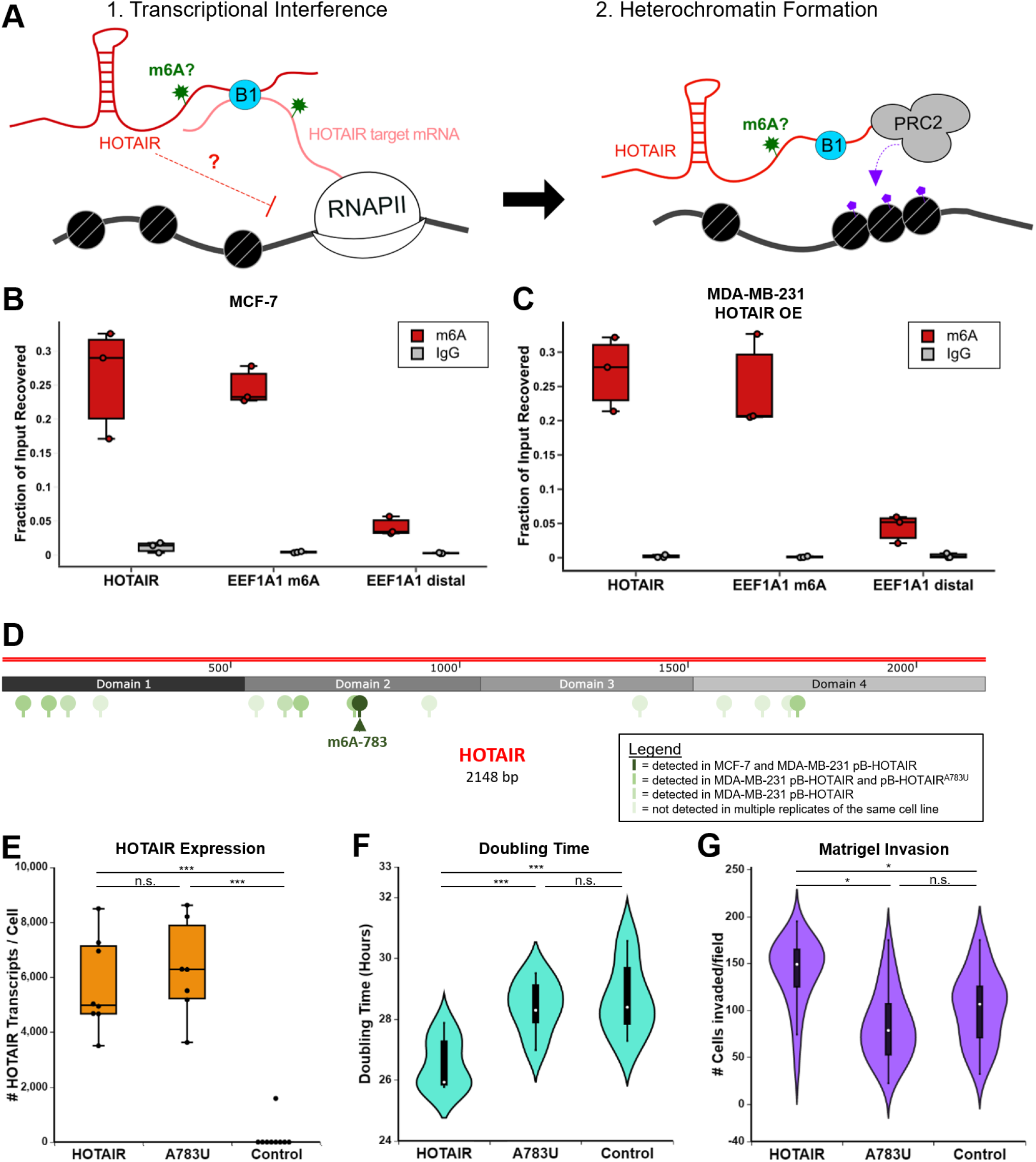
LncRNA HOTAIR is m6A modified. **A)** General model for HOTAIR mechanism. HOTAIR is initially recruited to its target loci via RNA-RNA interactions with its mRNA targets which is mediated by hnRNP B1. HOTAIR association with chromatin induces transcriptional interference via an unknown mechanism, promoting heterochromatin formation by PRC2 through H3K27me3. This paper investigates the role of m6A on HOTAIR. **B-C)** m6A RNA immunoprecipitation performed with an m6A antibody or IgG control in MCF-7 breast cancer cells (C) or MDA-MB-231 breast cancer cells with transgenic overexpression of HOTAIR (D). An m6A modified region in EEF1A1 (EEF1A1 m6A) is a positive control, while a distal region in EEF1A1 that is not m6A modified (EEF1A1 distal) serves as a negative control. **D)** m6A sites detected in HOTAIR-expressing cells in 6 experiments (yellow to red scale of increasing occurrences). m6A site 783 (dark red, arrow) was detected in every experiment except where it was mutated. **E)** Number of HOTAIR transcripts in MDA-MB-231 cells overexpressing WT HOTAIR, A783U HOTAIR, or an Anti-Luciferase control RNA. **F)** Doubling time of MDA-MB-231 overexpression cell lines described in (A). **G)** Quantification of cell invasion assays.

HOTAIR also interacts with lysine-specific demethylase 1 (LSD1), a histone demethylase that acts on H3K4me2, which has been proposed to reinforce repression by HOTAIR(L. Li et al., 2013; Somarowthu et al., 2015; Tsai et al., 2010). A new study in human epithelial kidney cells found that HOTAIR utilizes its LSD1-interacting domain to perturb LSD1 genomic distribution, independent of major changes in H3K4me2, leading to increased invasion(Jarroux et al., 2021). In this context, HOTAIR is proposed to inhibit the normal function of LSD1 in maintaining epithelial cells(Jarroux et al., 2021; McDonald, Wu, Timp, Doi, & Feinberg, 2011; Wang et al., 2009). In light of these findings, how HOTAIR specifically accomplishes transcriptional repression at its target loci, and how other pathways and cancer contexts influence HOTAIR function, remain elusive.

N6-methyladenosine (m6A) is a reversible RNA modification. It has been well studied in messenger RNAs (mRNAs), where it can regulate multiple steps of the mRNA life cycle, including processing, decay, and translation(Shi, Wei, & He, 2019); however, how m6A regulates lncRNA-mediated processes is less understood. Nevertheless, there is evidence for m6A regulation of lncRNAs. For example, the lncRNA Xist, a key mediator of X chromosome inactivation, contains multiple m6A sites that contribute to its ability to induce repression of the X chromosome(Coker et al., 2020; Patil et al., 2016).

The m6A modification on an RNA is typically recognized by a “reader” protein that binds specifically to methylated adenosine to mediate the functional outcome of m6A deposition. Apart from the YTH family of proteins which contain the YTH domain that directly read m6A, a handful of non-canonical indirect m6A readers have been suggested(Zaccara, Ries, & Jaffrey, 2019). In the case of Xist, the canonical YTH-containing nuclear localized m6A reader YTHDC1 recognizes m6A on Xist to mediate repression of the X chromosome(Nesterova et al., 2019; Patil et al., 2016). In contrast, m6A on *cis*-acting chromatin-associated regulatory RNAs leads to their YTHDC1-dependent degradation, preventing transcription of downstream genes(Jun Liu et al., 2020). Collectively, m6A influences the regulatory roles of both mRNA and noncoding RNA via diverse mechanisms(Coker, Wei, & Brockdorff, 2019).

RNA modifications such as m6A have been shown to play critical roles in several human cancers(X. Wu, Sang, & Gong, 2018). In breast cancer, studies have revealed that dysregulation of m6A levels can generate breast cancer stem-like cells and promote metastasis(Niu et al., 2019; C. Zhang et al., 2016; J. X. Zhang et al., 2013). Of the currently designated m6A reader proteins, we have previously shown that hnRNP A2/B1, a proposed non-canonical reader lacking the m6A-binding YTH domain, can interact with HOTAIR to regulate its chromatin and cancer biology mechanisms by promoting HOTAIR interactions with target mRNAs(Meredith et al., 2016). This evidence suggests that m6A may play a role in cancers where HOTAIR is overexpressed.

Here, we set out to investigate the potential function of m6A in HOTAIR-mediated breast cancer growth and invasion. We identify at least 8 m6A sites in HOTAIR and show that a single site (A783) is required for HOTAIR-mediated TNBC growth and invasion. Mutation of adenosine 783 in HOTAIR to uracil prevents the normal chromatin association and gene expression changes that are induced by the wild-type lncRNA. Surprisingly, the A783U mutant induces opposite gene expression changes to wild-type HOTAIR, reducing cancer phenotypes in TNBC cells, suggesting that the mutant HOTAIR is an antimorph. We find that YTHDC1, the nuclear m6A reader, interacts with HOTAIR at methylated A783 and artificial tethering of YTHDC1 at this site is sufficient to restore HOTAIR chromatin association in the A783 mutant. Finally, using a reporter system, we show that YTHDC1 mediates repression by HOTAIR in the absence of PRC2. Overall, our results suggest a model where a single site of m6A modification on HOTAIR enables a strong interaction with YTHDC1 to retain HOTAIR on chromatin for repression of its target genes, leading to altered TNBC properties. Collectively, our results demonstrate the potent activity of m6A on lncRNAs and in turn their role in cancer.

## Results

### HOTAIR contains multiple sites of m6A modification in breast cancer cell lines

To investigate the possibility that m6A regulates the function of HOTAIR in a mechanism similar to its regulation of lncRNA Xist, we examined previous genome-wide maps of m6A sites in human cells. Using the CVm6A database(Han et al., 2019), we found 3 m6A peaks in HOTAIR in HeLa cells, although the enrichment score for these sites was low(Figure 1 – figure supplement 1). To evaluate m6A methylation of HOTAIR in relevant breast cancer cells, we performed m6A RNA immunoprecipitation (meRIP) qRT-PCR in MCF-7 cells, which express low levels of endogenous HOTAIR(Meredith et al., 2016). A significant portion of HOTAIR was recovered upon immunoprecipitation with the anti-m6A antibody (26.2%, p=0.006), similar to an m6A modified region on the positive control region of *EEF1A1*, and consistently higher than a distal region of *EEF1A1* that is not m6A modified (Figure 1B).

We further found that m6A modification of HOTAIR is maintained during ectopic expression of HOTAIR in a stable MDA-MB-231 cell line. meRIP in MDA-MB-231 cells expressing transgenic HOTAIR resulted in significant HOTAIR recovery (27.1%, p=0.0009) (Figure 1C). These results demonstrate that HOTAIR is m6A modified in two distinct breast cancer contexts.

To identify single nucleotide sites of m6A modification, we performed a modified m6A eCLIP protocol(Roberts, Porman, & Johnson, 2020) on polyA-selected RNA from MCF-7 and MDA-MB-231 breast cancer cells (Figure 1 – figure supplement 2). In MCF-7 cells, we identified one m6A site within the HOTAIR transcript at adenosine 783 (Figure 1D, Table S1). m6A at adenosine 783 in MDA-MB-231 cells with transgenic HOTAIR was consistently detected with high confidence (Table S1), along with 7 other sites using our multi-replicate consensus approach (Roberts et al., 2020) (Table S2). Of note, A783 occurred within a non-canonical ‘GAACG’ sequence located in an unstructured region of the HOTAIR secondary structure(Somarowthu et al., 2015) (Figure 1 – figure supplement 3A).

To test if HOTAIR is m6A modified by the canonical m6A methyltransferase METTL3/14 complex, we performed shRNA mediated depletion of METTL3, METTL14, and the adaptor protein WTAP in MCF-7 cells (Figure 1 – figure supplement 4B). We observed a ~3 to 5-fold reduced recovery of HOTAIR in methyltransferase-depleted cells relative to non-targeting controls (p=0.0063) (Figure 1 – figure supplement 4C). Together, these results indicate that m6A methylation of HOTAIR is dependent on the METTL3/14 complex.

### Nucleotide A783 is important for the ability of HOTAIR to promote breast cancer cell proliferation and invasion

Given that nucleotide A783 was consistently methylated within HOTAIR in our m6A mapping experiments, in both endogenous and overexpressed contexts, we asked whether this modification had any consequences to HOTAIR function. To directly test the functional role of A783, we mutated the adenosine to uracil at this position (HOTAIR^A783U^). We then mapped m6A sites in MDA-MB-231 cells overexpressing the HOTAIR^A783U^ mutant as above (Table S1). Both wild-type (WT) and the mutant form of HOTAIR were expressed at similar levels, with approximately 5,000 transcripts per cell (Figure 1E), resembling the high levels of HOTAIR observed in samples from cancer patients(Arshi A, Raeisi F, Mahmoudi E, Mohajerani F, Kabiri H, Fazel R, Zabihian-Langeroudi M, 2020; Gupta et al., 2010; Yang et al., 2011). While the CLIP-based m6A signature was no longer detected at adenosine 783 when this site was mutated to uracil, we detected m6A modification at five of the seven other multi-replicate consensus sites (Tables S1 and S2, Figure 1B). Nucleotides 143 and 620 were no longer called with multi-replicate consensus confidence as m6A in the A783U mutant, though m6A143 was only called in WT HOTAIR at our lowest confidence category and m6A620 is called in one of the A783U mutant replicates (Table S1). Nonetheless, it is possible that methylation at A783 is required for one or both m6A events to occur.

To determine the effect of the A783U mutation on HOTAIR-mediated breast cancer cell growth, we measured the doubling time of MDA-MB-231 cells expressing WT and A783U mutant HOTAIR. As described above, we overexpressed HOTAIR and the HOTAIR^A783U^ mutant in MDA-MB-231 cells and included overexpression of an antisense sequence of luciferase mRNA (Anti-Luc) as a negative control(Meredith et al., 2016). Similar to previous studies, transgenic overexpression of HOTAIR resulted in 10^3^-10^4^ copies of HOTAIR per cell and mediated increased cancer growth and invasion of MDA-MB-231 cells(Gupta et al., 2010) (Figure 1E-G). We performed cell proliferation assays by plating 5,000 cells in a 96-well dish and analyzing confluency every 2 hours over a period of 48 hours (example shown in Figure 1 – figure supplement 3B). We observed that MDA-MB-231 cells overexpressing WT HOTAIR proliferated more quickly, with a shorter doubling time (~26 hours) than cells overexpressing Anti-Luc (~28.5 hours, p=0.0003) (Figure 1F and Figure 1 – figure supplement 3C). Surprisingly, the single nucleotide mutation of A783U in HOTAIR abolished its ability to enhance MDA-MB-231 cell proliferation; cells expressing HOTAIR^A783U^ proliferated more slowly, with a longer doubling time than those expressing WT HOTAIR (~28.6 hours, p=0.004) and grew similarly to cells containing the Anti-Luc control. To examine the role of A783 of HOTAIR in mediating breast cancer cell invasion, the same MDA-MB-231 cell lines were plated in a Matrigel invasion assay. Overexpression of WT HOTAIR induced a significant increase in number of cells invaded compared to the Anti-Luc control (p=0.038). In contrast, overexpression of A783U HOTAIR did not lead to an increase in invasion compared to the Anti-Luc control (p=0.22) and resulted in significantly less cells invaded compared to overexpression of WT HOTAIR (p=0.012) (Figure 1G). Altogether, these results suggest that m6A modification of adenosine 783 in HOTAIR is key for mediating the increased aggressiveness of TNBC that is promoted in contexts where the lncRNA is overexpressed.

### Overexpression of A783U mutant HOTAIR induces divergent gene expression changes from wild-type HOTAIR in breast cancer cells

To analyze HOTAIR-mediated gene expression changes in MDA-MB-231 cells, we performed high throughput RNA sequencing on cells overexpressing WT HOTAIR, A783U mutant HOTAIR, or antisense luciferase as a control. For cells expressing WT HOTAIR, we identified 155 genes that were differentially expressed (adjusted p<0.1) when compared with control cells expressing Anti-Luciferase (Figure 2A). Upregulated genes in cells expressing WT HOTAIR include genes involved in positive regulation of angiogenesis (p=1.22E-05), regulation of cell population proliferation (p=0.0361), and cell differentiation (p=0.0157), while downregulated genes include genes involved in cell adhesion (p=0.0118), p53 (p=0.0112) and MAPK (p=0.0313) signaling, and tumor repressors such as HIC1 and DMNT3A. This set of genes had significantly different expression in cells overexpressing WT HOTAIR compared to either cells overexpressing Anti-Luciferase or the A783U mutant HOTAIR (Figure 2A-C). Surprisingly, mutation of A783 did not merely prevent most gene expression changes seen in wild-type HOTAIR, but instead, expression of the A783U mutant induced certain changes in the opposite direction from the baseline control MDA-MB-231 cell line. We confirmed this pattern by qRT-PCR: genes that were upregulated in MDA-MB-231 cells upon introduction of WT HOTAIR had decreased expression in cells with A783U mutant HOTAIR (Figure 2B). This included genes such as *PTK7* involved in the Wnt signaling pathway (fold change relative to control in WT HOTAIR=2.6, p=0.008; in A783U HOTAIR=−1.4, p=0.002); *CDH11*, a mesenchymal cadherin that is upregulated in invasive breast cancer cell lines(Pishvaian et al., 1999) (fold change in WT HOTAIR=2.0, p=0.01; in A783U HOTAIR=−1.6, p=0.02); and *GRIN2A*, an oncogenic glutamate receptor (fold change in WT HOTAIR=2.5, p=0.006; in A783U HOTAIR=−3.7, p=9.3E-05). Similarly, genes downregulated with WT HOTAIR were significantly increased with A783U HOTAIR, compared to the parental MDA-MB-231 control, including *SEMA5A*, a guidance cue protein that suppresses the proliferation and migration of lung adenocarcinoma cells(Ko et al., 2020) (fold change relative to control in WT HOTAIR=−3.5, p=0.0002; in A783U HOTAIR=2.8, p=0.01); *SIRPA*, a cell surface receptor that can act as a negative regulator of the phosphatidylinositol 3-kinase signaling and mitogen-activated protein kinase pathways(Takahashi, 2018) (fold change in WT HOTAIR=−1.6, p=0.03; in A783U HOTAIR=3.1, p=0.009); and *TP53I11*, a p53-interacting protein that suppresses migration and metastasis in MDA-MB-231 cells(Xiao et al., 2019) (fold change in WT HOTAIR=-1.7, p=0.03; in A783U HOTAIR=4.0, p=0.008) (Figure 2C). To further analyze differences in cells expressing A783U mutant HOTAIR, we performed a pairwise comparison with control MDA-MB-231 cells and identified 758 differentially expressed genes (Figure 2D). Upregulated gene categories in A783U HOTAIR-expressing cells include negative regulation of response to growth factor stimulus (p=2.27E-04), positive regulation of apoptosis (p=3.88E-04), and regulation of migration (p=4.36E-04), while downregulated gene categories include regulation of the epithelial to mesenchymal transition (p=1.48E-04), angiogenesis (p=1.64E-04), cell adhesion (p=7.51E-05), and cell migration (p=1.64E-04). We hypothesize that this altered pattern of gene expression may underlie the slight decrease in cell invasion observed in the A783U context compared to control MDA-MB-231 cells (Figure 1G).

**Figure 2.**
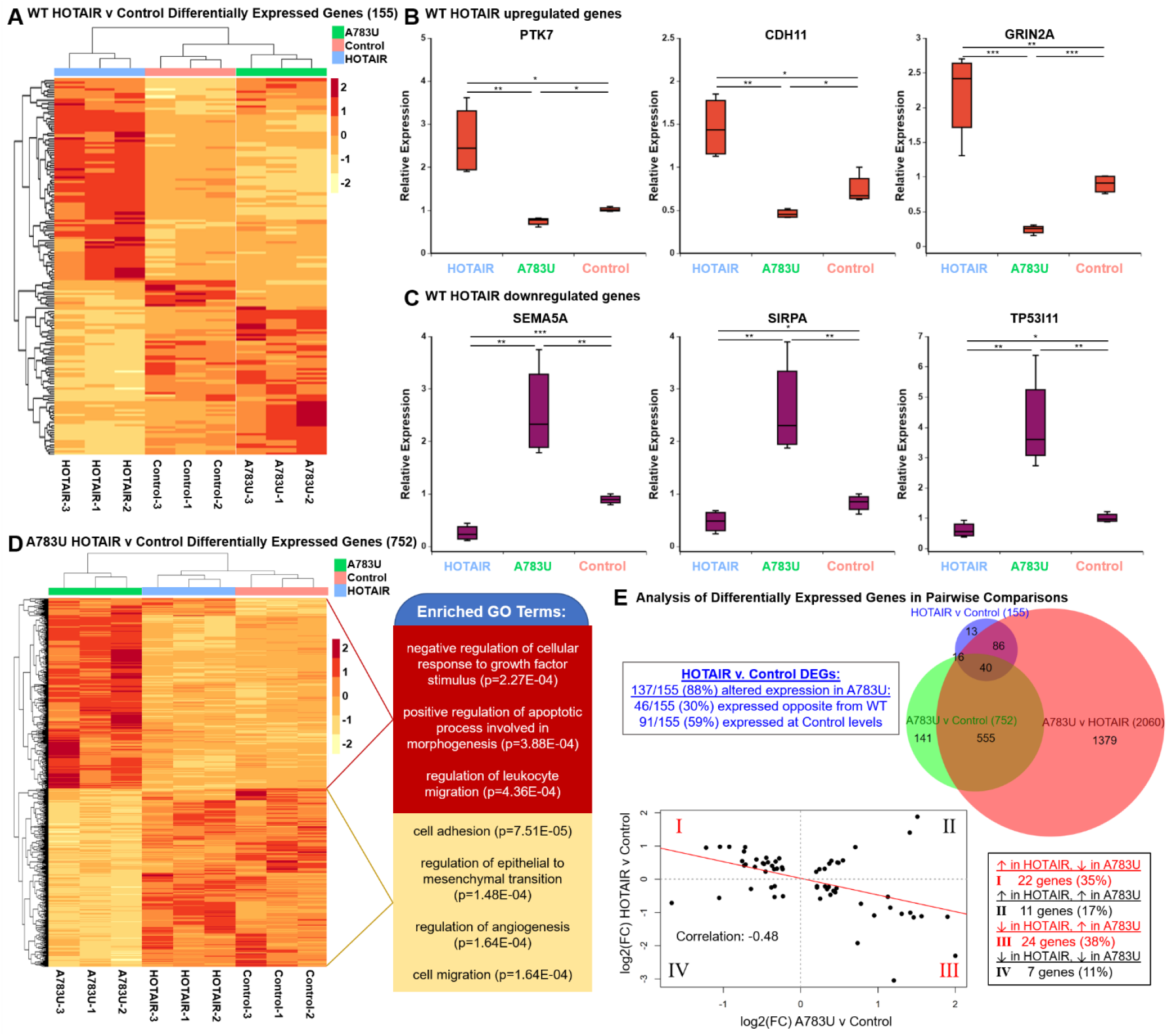
HOTAIR-mediated gene expression changes in breast cancer are altered by mutation of A783. **A)** Heatmap of Z-scores of differentially expressed genes (DEGs) between MDA-MB-231 cells overexpressing wild-type HOTAIR versus an Anti-Luciferase control. **B-C)** qRT-PCR analysis of genes upregulated (B) or downregulated (C) upon HOTAIR overexpression. **D)** Heatmap of Z-scores of DEGs between MDA-MB-231 cells overexpressing A783U mutant HOTAIR versus an Anti-Luciferase control, left. Selected significant GO terms in upregulated (red) and downregulated (yellow) genes, right. **E)** Additional analysis of differentially expressed genes. Top, Venn diagram (created using BioVenn, (Hulsen, de Vlieg, & Alkema, 2008)) of number of DEGs between MDA-MB-231 cells overexpressing wild-type HOTAIR, A783U mutant HOTAIR, or an Anti-Luciferase control. Top left inset describes the direction of change in the A783U v. Control relative to the direction of the wild-type HOTAIR v. Control, based on adjusted p<0.1. Bottom, correlation analysis of expression in HOTAIR v Control and A783U v Control pairwise comparisons. Linear regression was used to fit a trend line (red) over the points, with the calculated Pearson correlation coefficient included in the graph. Bottom right inset describes the number of genes in each quadrant.

To further investigate differences between cells expressing WT HOTAIR versus the A783U mutant HOTAIR, we performed a pairwise comparison. Here, we observed the most differentially expressed genes (2060) compared to other pairwise comparisons (Figure 2E, top). Overall, these results reveal that expression of the A783U mutant HOTAIR induces additional and often opposite gene expression changes compared to expression of WT HOTAIR in breast cancer cells, suggesting a potential antimorph property of this single nucleotide mutation. The opposite gene expression pattern is evident in the heat map of all differentially expressed genes (Figure 2 – figure Supplement 1A), as well as the observation that most (137/155, 88%) of WT HOTAIR-regulated genes have altered expression with A783U HOTAIR, with a significant portion (46/155, 30%) having opposite expression in MDA-MB-231 cells expressing A783U HOTAIR compared to control cells (Figure 2E, Figure 2 – figure Supplement 1B-C). This pattern is also evident in the negative correlation (−0.48) when fold change in expression for HOTAIR v Control and A783U v Control is plotted (Figure 2E, bottom). We hypothesized that prevention of m6A methylation by the A783U mutation disrupts an m6A-dependent function to cause loss-of-function and antimorph cell biology and gene expression behaviors.

### hnRNP B1 is not a direct m6A reader in MCF-7 cells

We next sought to address the mechanisms behind HOTAIR m6A783 function. hnRNP A2/B1 has previously been suggested to be a reader of m6A, and the B1 isoform has a high affinity for binding HOTAIR(Alarcon et al., 2015; Meredith et al., 2016; Yingmin Wu et al., 2019). However, comparing our previously generated eCLIP results for hnRNP B1(Nguyen, Balas, Griffin, Roberts, & Johnson, 2018) to the m6A eCLIP, both performed in MCF-7 cells, we found that, out of 10,470 m6A sites, only 417 (4%) were identified to contain an hnRNP B1 binding site within 1,000 nucleotides (Figure 3 – figure supplement 1A). Upon mapping hnRNP B1 signal intensity relative to the nearby m6A site, we observed that hnRNP B1 is depleted directly over m6A sites (Figure 3 – figure supplement 1B). These results suggest that hnRNP B1 is not a direct m6A reader, although m6A may indirectly promote its recruitment in some contexts. When comparing hnRNP B1 binding in HOTAIR with m6A sites, B1 binding peaks in MCF-7 cells occur in m6A-free regions of HOTAIR. Conversely, data from *in vitro* eCLIP analysis of B1 binding to unmodified HOTAIR reveal additional B1 binding peaks in Domain 1 of HOTAIR, one of which occurs near several m6A sites (Figure 3 – figure supplement 1C). Altogether, these data suggest that m6A is not likely to directly recruit hnRNP B1 as a reader, although it could contribute to hnRNP B1 binding.

**Figure 3.**
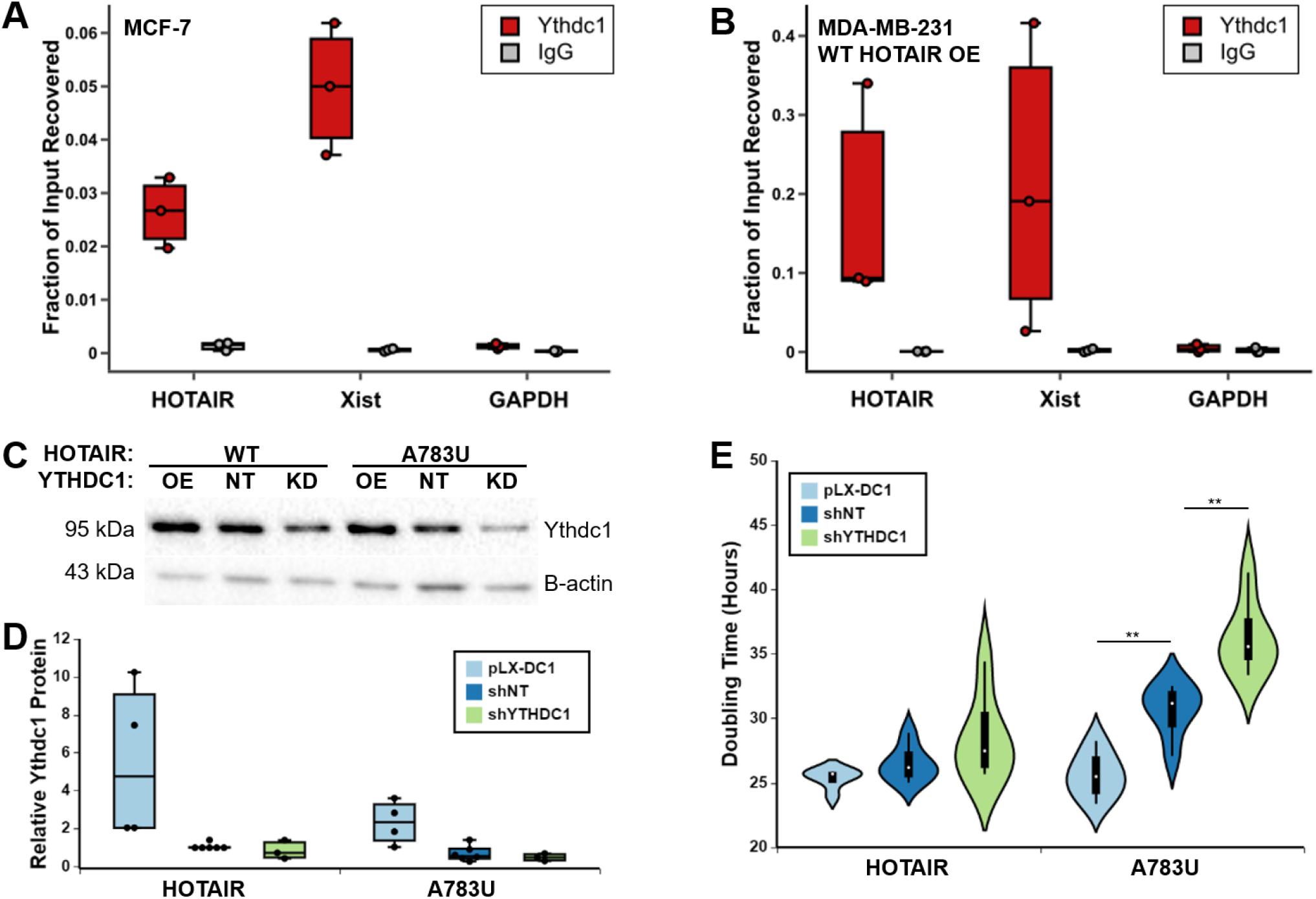
YTHDC1 interacts with HOTAIR and enables HOTAIR-mediated breast cancer growth. **A-B)** YTHDC1 RIP performed in MCF-7 cells (A) or MDA-MB-231 cells overexpressing transgenic HOTAIR (B). **C)** Western blot results of YTHDC1 protein levels in pLX-DC1 overexpression, shNT control, and shDC1 knockdown MDA-MB-231 cell lines expressing WT or A783U HOTAIR. **D)** Quantification of 3 replicates of (C). Protein levels of YTHDC1 were normalized to β-actin levels and are relative to the HOTAIR shNT sample. **E)** Doubling time of MDA-MB-231 cells containing WT or A783U HOTAIR and overexpression or knockdown of YTHDC1.

### YTHDC1 interacts with HOTAIR to mediate breast cancer proliferation

In light of the results for hnRNP A2B1 described above, we turned to alternative candidate m6A readers of HOTAIR. YTHDC1 is a nuclear-localized m6A reader that binds m6A sites in noncoding RNAs, including the Xist lncRNA (Patil et al., 2016). We reasoned that YTHDC1 was a strong candidate for interaction with HOTAIR, which is a lncRNA that is also primarily nuclear-localized. To determine if YTHDC1 interacts with HOTAIR, we performed RNA immunoprecipitation (RIP) qRT-PCR using an antibody to YTHDC1. In both MDA-MB-231 cells overexpressing transgenic HOTAIR, and MCF-7 cells expressing endogenous HOTAIR, a significant portion of HOTAIR RNA was recovered when using antibodies specific against YTHDC1 (17.4%, p=0.04; 2.6%, p=0.003, respectively) (Figure 3A-B).

To test the role of YTHDC1 in HOTAIR’s ability to enhance breast cancer cell proliferation, we stably overexpressed or knocked down YTHDC1 in the context of WT or A783U HOTAIR overexpression in MDA-MB-231 cells (Figure 3C-D). We noted that YTHDC1 protein levels tended to be ~2-fold higher in cells containing WT HOTAIR compared to A783U mutant HOTAIR (Figure 3D). Although this difference was not significant (p=0.16), it suggests a potential positive relationship between WT HOTAIR RNA and YTHDC1 protein levels. Next, we used the MDA-MB-231 cell lines we generated to analyze proliferation as described above. Growth of MDA-MB-231 cells overexpressing WT HOTAIR was not significantly altered by YTHDC1 dosage (0.96 fold change, p=0.16 for pLX-DC1; 1.08 fold change, p=0.26 for shDC1, respectively), yet there was a trend towards decreased doubling time with increasing YTHDC1. In contrast, cells with A783U mutant HOTAIR had significant differences in doubling time with overexpression or knockdown of YTHDC1 (Figure 3E). Overexpression of YTHDC1 led to significantly faster growth of MDA-MB-231 cells containing A783U mutant HOTAIR (0.84-fold change in doubling time, p=0.003), with proliferation rates comparable to cells expressing WT HOTAIR. Knockdown of YTHDC1 in cells containing HOTAIR^A783U^ was particularly potent in reducing the growth rate (~1.2-fold increase in doubling time, p=0.008) demonstrating a role for YTHDC1 in mediating HOTAIR’s ability to enhance proliferation of breast cancer cells through A783, or via other m6A sites in A783U mutant HOTAIR upon YTHDC1 overexpression.

### High *HOTAIR* levels are associated with an aggressive disease progression in breast cancer patients with high tumor *YTHDC1* expression

To further explore a clinical role for HOTAIR and YTHDC1 in breast cancer, we used GEPIA2, a web server for large-scale expression profiling and interactive analysis (Tang, Kang, Li, Chen, & Zhang, 2019). To this end, we analyzed the relationship of *HOTAIR* and *YTHDC1* expression in publicly available outcomes data from breast cancer patient primary tumors. Across all breast cancer patient samples, *YTHDC1* is generally expressed at the mRNA level, ranging roughly five-fold. HOTAIR levels vary more widely, with some samples not expressing the lncRNA. Because of these different expression profiles, there is only a very modest positive correlation between *HOTAIR* and *YTHDC1* (R=0.092, p=0.0025) (Figure 4 – figure supplement 1A). To further investigate *HOTAIR* and *YTHDC*1 in breast tumors, we used the Kaplan-Meier Plotter to analyze recurrence-free and overall survival of breast cancer patients(Gyorffy et al., 2010) as well as UALCAN (a tool for analyzing cancer OMICS data) to determine gene expression in normal breast tissue versus breast tumor specimens(Chandrashekar et al., 2017). Consistent with previous studies, high expression of *HOTAIR* is indicative of a shorter time to recurrence (HR=1.41, p=6.3e-05) (Figure 4A) and shorter overall survival (HR=1.65, p=0.0084) (Figure 4 – figure supplement 1B)(Arshi A, Raeisi F, Mahmoudi E, Mohajerani F, Kabiri H, Fazel R, Zabihian-Langeroudi M, 2020; Gupta et al., 2010). *HOTAIR* RNA expression is increased in breast cancer specimens ~7-fold compared to normal breast tissues (p=1.62e-12), with the highest *HOTAIR* expression (~14.5-fold increase) observed in Stage 4 disease (p=5.36e-04) (Figure 4 – figure supplement 1C-D). The reverse is true for *YTHDC1*, with high levels corresponding to longer disease-free status (HR=0.69, p=2.5e-11) (Figure 4B) and overall survival (HR=0.73, p=0.0088) (Figure 4 – figure supplement 2A) and a modest decrease (~10%) in mRNA levels in tumor compared to normal tissue (p=1.14e-06) (Figure 4C, Figure 4 – figure supplement 2B). Interestingly, YTHDC1 protein is higher in tumor samples compared to normal tissue (p=6.7e-09) (Figure 4D, Figure 4 – figure supplement 2C). This could be because in general there are fewer epithelial cells in normal breast compared to the number of carcinoma cells in breast tumors(Rezaul et al., 2010), and may suggest a significant amount of translational regulation for the YTHDC1 mRNA.

**Figure 4.**
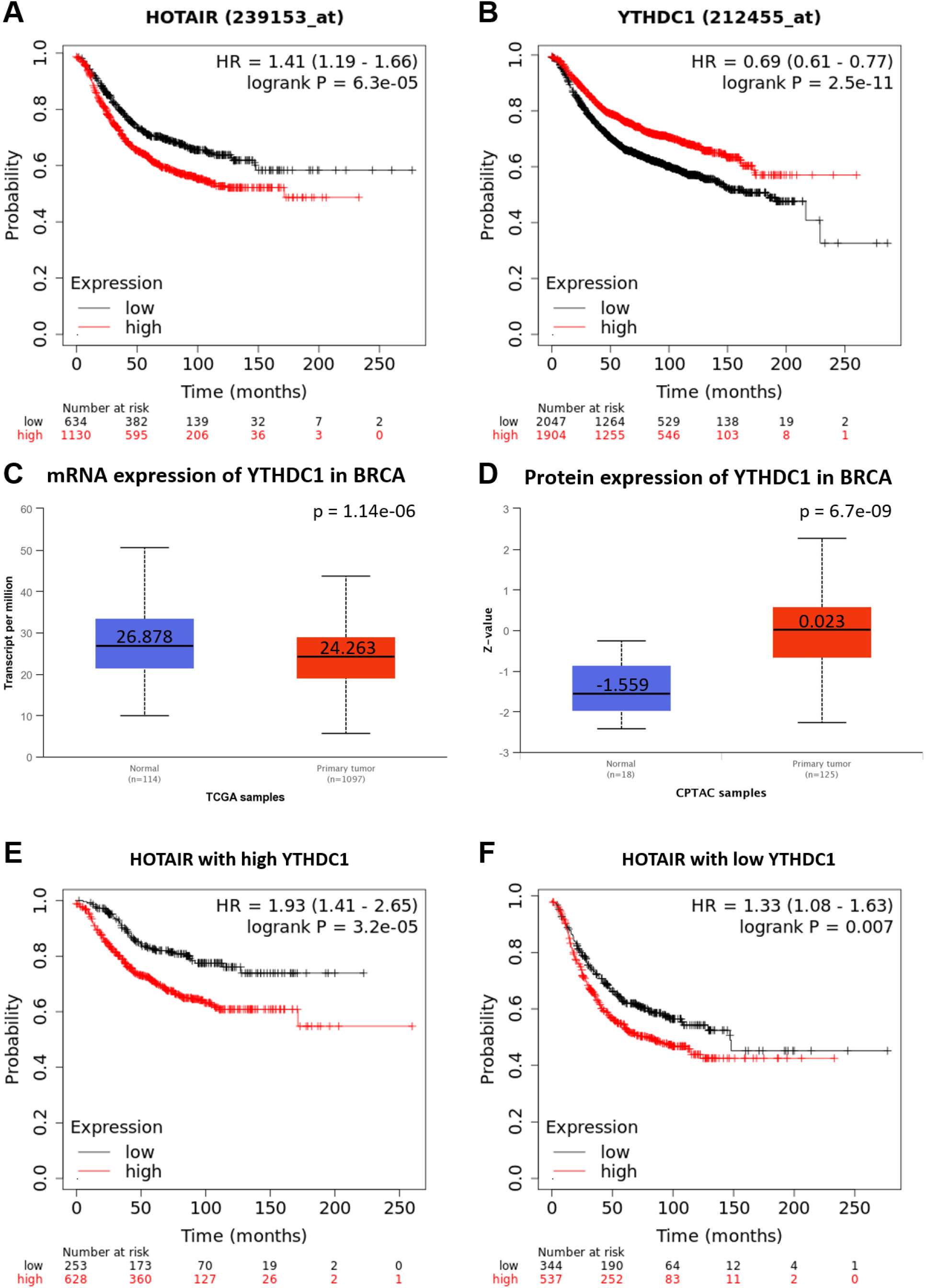
*YTHDC1* and *HOTAIR* in breast cancer outcomes. **A-B)** Kaplan-Meier curves for recurrence-free survival of breast cancer patients with high or low expression of A) *YTHDC1* or B) *HOTAIR* generated using Kaplan-Meier Plotter(Gyorffy et al., 2010). **C-D)** Expression of YTHDC1 C) mRNA and D) protein in normal breast tissue versus breast cancers generated with UALCAN(Chandrashekar et al., 2017). **E-F)** Recurrence-free survival curves for breast cancer patients examining effect of HOTAIR on the background of either E) high or F) low *YTHDC1* levels, generated with Kaplan-Meier Plotter(Gyorffy et al., 2010).

To examine *YTHDC1* in relation to *HOTAIR* in breast cancer outcomes, we assessed recurrence-free and overall survival based on *HOTAIR* expression in cohorts of tumors expressing either high or low levels of *YTHDC1* mRNA. In the context of high *YTHDC1* expression, *HOTAIR* is even more strongly indicative of risk for shorter time to recurrence (HR=1.93, p=3.2e-05) (Figure 4E) and shorter overall survival (HR=2.1, p=0.0012) (Figure 4 – figure supplement 2D) compared to *HOTAIR* alone (Figures 4B, S6B). On the background of low *YTHDC1*, *HOTAIR* has a less impressive effect on disease-free survival (HR=1.33, p=0.007) (Figure 4F), more similar to the effect of *HOTAIR* alone (Figure 4B), and on overall survival, where *HOTAIR* expression is no longer a significant prognostic indicator (HR=1.34, p=0.23) (Figure 4 – figure supplement 2E). Altogether, these patient outcomes data are consistent with high *YTHDC1* levels potentially contributing to the ability of HOTAIR to affect breast cancer progression.

### Mutation at A783 of HOTAIR results in decreased interaction with YTHDC1 *in vitro*, but does not abolish HOTAIR methylation or YTHDC1 interaction at other sites *in vivo*

To determine if nucleotide A783 in HOTAIR recruits YTHDC1 via m6A modification, we generated PP7-tagged *in vitro* transcribed RNA of domain 2 of WT or A783U mutant HOTAIR and performed *in vitro* m6A methylation with purified METTL3/14(Jianzhao Liu et al., 2014) and S-adenosylmethionine as a methyl donor. We then transfected HEK293 cells with an expression plasmid containing FLAG-tagged YTHDC1 and obtained protein lysates. The *in vitro HOTAIR* transcripts were tethered to IgG-coupled magnetic beads via a PP7-Protein A fusion protein and incubated with FLAG-YTHDC1-containing protein lysates. Beads were washed and the relative recovery of FLAG-YTHDC1 was determined by anti-FLAG Western Blot (Figure 5A). WT HOTAIR interaction with YTHDC1 was enhanced when the transcript was m6A-modified (~3-fold increase, p=0.04), while A783U HOTAIR interaction with YTHDC1 was not significantly altered by the addition of m6A (~1.3-fold change, p=0.6) (Figure 5B-C).

**Figure 5.**
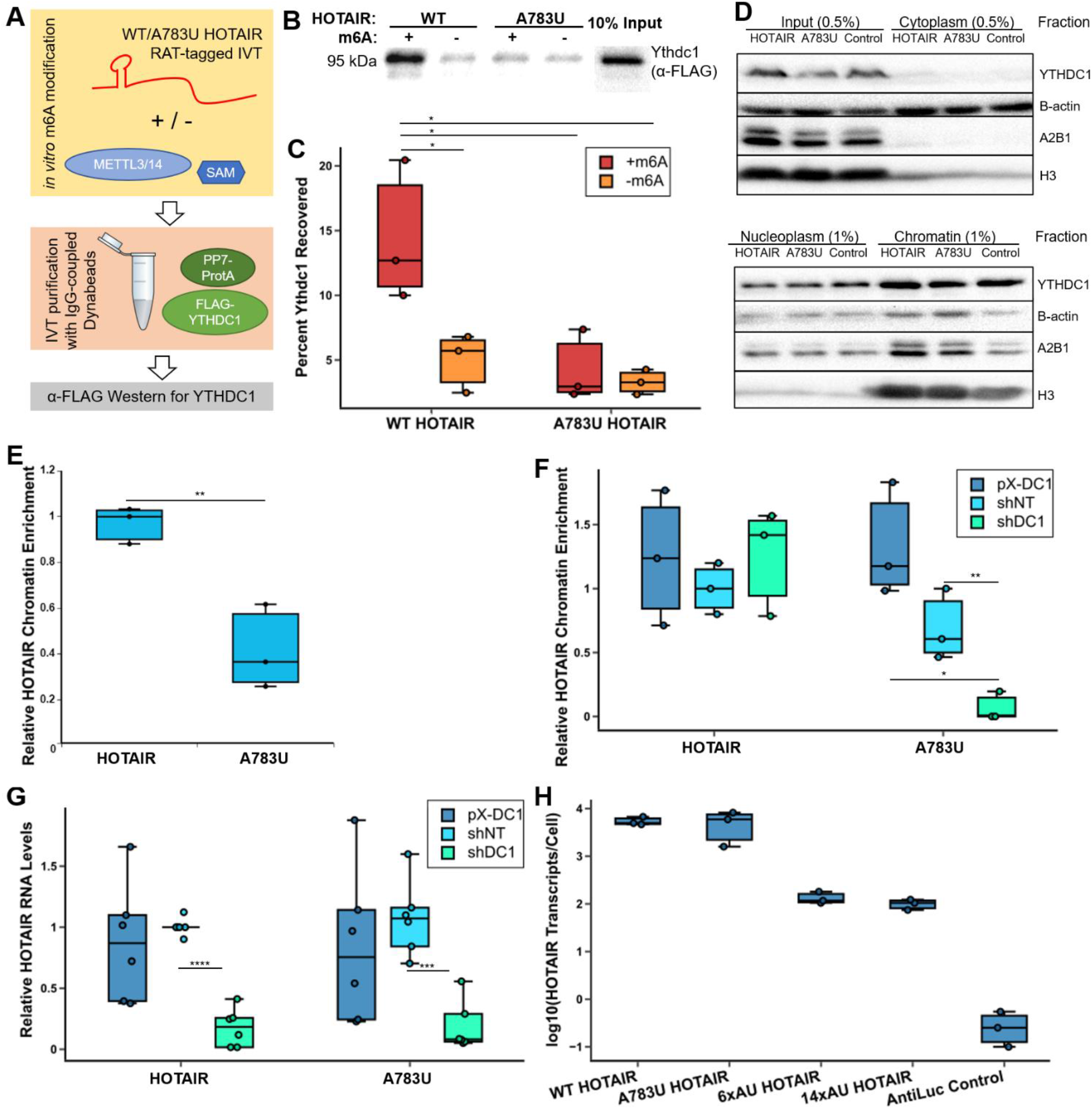
HOTAIR m6A site 783 mediates interaction with YTHDC1 and chromatin association. **A)** Schematic of YTHDC1 pulldown experiment with m6A-modified WT and A783U HOTAIR. PP7-tagged domain 2 of WT or A783U HOTAIR was *in vitro* transcribed and m6A modified with purified METTL3/14. RNA was bound to PP7-Protein A, cellular extract containing FLAG-tagged YTHDC1 was added, and a pulldown was performed with IgG-coupled Dynabeads. Amount of FLAG-YTHDC1 bound was assessed by Western blot. **B)** Anti-FLAG Western blot of pulldown experiment outlined in (A). **C)** Quantification of anti-FLAG Western blots from 3 replicates. **D)** Western blot performed on fractionation of MDA-MB-231 cell lines overexpressing WT or A783U HOTAIR or Antisense-Luciferase. **E)** qRT-PCR was performed on fractionated RNA samples from MDA-MB-231 cells containing overexpression of WT or A783U HOTAIR, and chromatin association was calculated by determining the relative chromatin-associated RNA to input and normalizing to 7SL levels and relative to WT HOTAIR samples. **F)** Chromatin enrichment was calculated similarly as in (D) in MDA-MB-231 cell lines expressing WT or A783U HOTAIR with knockdown or overexpression of YTHDC1. Values are relative to HOTAIR shNT samples. **G)** qRT-PCR of *HOTAIR* RNA levels in MDA-MB-231 cell lines overexpressing WT or A783U HOTAIR containing overexpression or knockdown of YTHDC1. **H)** qRT-PCR of *HOTAIR* RNA levels in MDA-MB-231 cell lines expressing WT, A783U, 6xAU, or 14xAU HOTAIR or an AntiLuc control.

To characterize changes to the molecular interactions that occur with mutation of A783 in breast cancer cells, we performed m6A and YTHDC1 RIP experiments on MDA-MB-231 cells overexpressing WT or A783U HOTAIR (Figure 5 – figure supplement 1A-B). Surprisingly, we did not see any significant changes in HOTAIR recovery in either experiment (~1.2-fold change, p=0.5; 1.1-fold change, p=0.8 for m6A and YTHDC1 RIP, respectively). The HOTAIR^A783U^ maintains m6A modifications at other sites within the RNA, which we have mapped in the overexpression context (Table S1). These sites are likely sufficient for HOTAIR recovery when immunoprecipitating YTHDC1. However, modification of A783, in particular, appears to be important in mediating the physiological effects observed by HOTAIR overexpression, likely by YTHDC1 binding to A783 in a methylation-dependent manner, as observed in the *in vitro* experiment (Figure 5B-C).

### m6A and YTHDC1 mediate chromatin association and expression of HOTAIR

Based on the differences observed between cell lines containing WT and A783U HOTAIR and the function of HOTAIR in chromatin-mediated gene repression, we investigated whether chromatin association of HOTAIR was altered in these cells. We performed fractionation of MDA-MB-231 cells containing WT or A783U HOTAIR or an antisense-Luciferase control into cytoplasm, nucleoplasm, and chromatin fractions (Figure 5D, see Methods). We isolated RNA from each fraction and performed qRT-PCR for HOTAIR and GAPDH. Cells overexpressing WT HOTAIR had significantly more chromatin-associated HOTAIR (~4.3-fold) than cells expressing A783U HOTAIR (p<0.05) (Figure 5E), though overall levels of HOTAIR are unchanged (Figure 2A).

To examine the effect of YTHDC1 levels on HOTAIR chromatin association, we performed a similar fractionation experiment in MDA-MB-231 cells expressing WT or A783U HOTAIR with overexpression or knockdown of YTHDC1 (Figure 5 – figure supplement 1C). While YTHDC1 levels did not significantly alter WT HOTAIR chromatin association, overexpression of YTHDC1 increased HOTAIR^A783U^ chromatin association ~1.9-fold to similar levels as WT HOTAIR (p=0.05), and knockdown resulted in a significant ~10-fold decrease in chromatin association (p=0.01) (Figure 5F). We reason that the differences observed between WT and A783U mutant HOTAIR are due to a high affinity constitutive interaction of YTHDC1 with WT HOTAIR at m6A783 that enables chromatin association and is not affected by knockdown or overexpression. For A783U mutant HOTAIR that does not interact with YTHDC1 at this position, increasing the concentration of YTHDC1 can drive interaction at other (lower affinity) m6A sites within the mutated HOTAIR. These interactions occur at a low level in cells with wild-type YTHDC1 levels, and, since they are low affinity, are most sensitive to knockdown of YTHDC1 (Figure 5 – figure Supplement 1D). Therefore, the A783U mutant, which only retains these proposed lower affinity sites, is particularly sensitive to YTHDC1 levels.

While HOTAIR expression levels remained similar for DC1 overexpression lines compared to shNT control lines (0.8 fold change, p=0.3), they were significantly decreased by ~5 to 10-fold in YTHDC1 knockdown lines for both WT and A783U mutant HOTAIR relative to shNT cell lines (p=3.24e-10) (Figure 5G). These results suggest that YTHDC1 regulates the expression or stability of HOTAIR, independently of A783. To investigate the role of other m6A sites within HOTAIR, we generated HOTAIR overexpression constructs containing 6 or 14 adenosine-to-uracil mutations (6xAU and 14xAU, respectively) both of which included A783U. While WT and A783U HOTAIR expression levels were similarly high, there was a ~50-fold decrease in expression of 6xAU or 14xAU HOTAIR (Figure 5H). This suggests that other m6A sites within HOTAIR mediate its high expression levels in breast cancer cells.

### Tethering YTHDC1 to A783U mutant HOTAIR restores chromatin association

To more directly examine the effects of YTHDC1 interaction with HOTAIR in the context of the A783U mutation, we employed a catalytically inactive RNA-targeting Cas protein, dCasRX, which has previously been used to recruit effectors to specific RNA molecules via a guide RNA (Figure 6A)(Konermann et al., 2018). We transfected MDA-MB-231 cells stably expressing WT or A783U HOTAIR with a plasmid containing the dCasRX-YTHDC1 fusion protein, in combination with a plasmid containing either a HOTAIR guide RNA (targeting a 22-nucleotide sequence 7 nucleotides downstream from A783 in HOTAIR, see Figure 6A) or a non-targeting gRNA. Expression of dCasRX-YTHDC1 was confirmed by Western blot (Figure 6B). While chromatin association of WT HOTAIR remained consistently high, chromatin association levels of A783U HOTAIR were only restored to near WT HOTAIR levels upon transfection with plasmids containing the dCasRX-YTHDC1 fusion protein and the HOTAIR gRNA (p=0.25 compared to WT HOTAIR). In contrast, chromatin association of A783U HOTAIR remained low upon transfection of dCasRX-YTHDC1 with a non-targeting guide RNA (~3.7 fold lower than WT HOTAIR, p=0.0066) (Figure 6C). HOTAIR RNA levels remained consistent in all samples (Figure 6D). These results confirm that YTHDC1 mediates chromatin localization of HOTAIR, and show that the chromatin association defect of the A783U mutation can be restored simply by restoring binding of YTHDC1 at that specific location.

**Figure 6.**
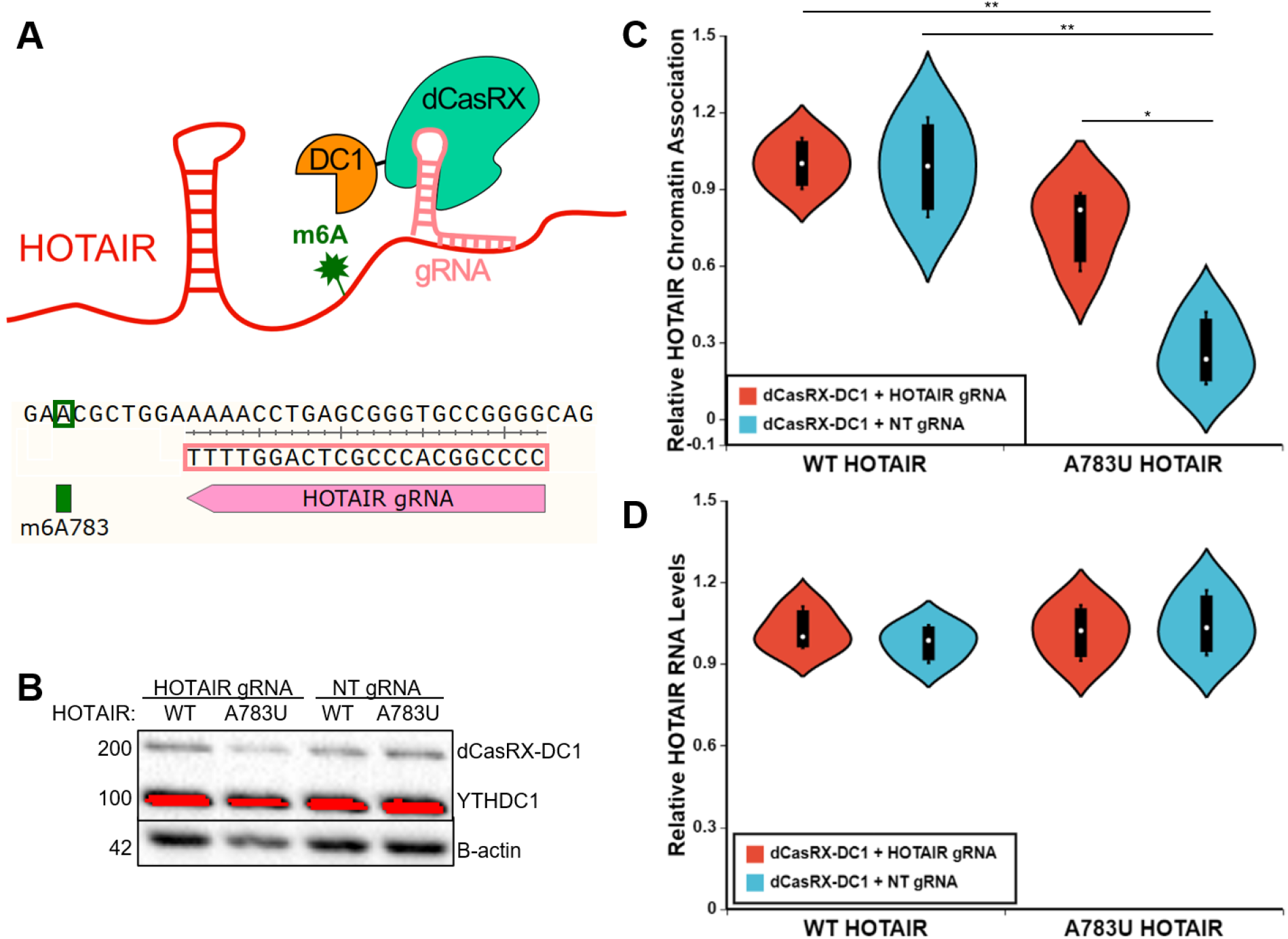
Tethering YTHDC1 to A783U mutant HOTAIR restores chromatin localization independent of changes in RNA levels. **A)** Schematic of tethering strategy using a dCasRX-YTHDC1 fusion protein and a guide RNA targeted just downstream of A783 in HOTAIR. **B)** Examples of Western blots for YTHDC1 (upper) and B-actin (lower) on input, cytoplasmic, nucleoplasmic, and chromatin samples, as noted. **C)** Similar analysis described in Figure 5D-E was performed on fractionated RNA samples from cell lines overexpressing WT or A783U HOTAIR transfected with a plasmid containing dCasRX-YTHDC1 in combination with a HOTAIR or non-targeting (NT) gRNA, as noted. **D)** Relative HOTAIR RNA levels in Input samples from C.

### YTHDC1 contributes to gene repression by HOTAIR in the absence of PRC2, independent of its role in chromatin association or RNA stability

To determine the effect of YTHDC1 on transcriptional repression mediated by HOTAIR, we used previously generated reporter cell lines that contain HOTAIR artificially directly tethered to chromatin upstream of a luciferase reporter gene to repress expression, independent of PRC2(Portoso et al., 2017) (Figure 7A). We confirmed that HOTAIR tethered upstream of the luciferase reporter reduced luciferase expression using both qRT-PCR (~2.3-fold lower, p=8.0e-12) and luciferase assay (~3.1-fold lower, p=0.002) (Figure 7B,C). We also performed m6A eCLIP to confirm that *HOTAIR* was m6A modified in this context and detected 10 m6A sites within *HOTAIR*, including A783 (Table S1). To test the role of YTHDC1 in the repression mediated by HOTAIR, we used 3 different siRNAs to knock down YTHDC1 relative to a non-targeting control (~2-fold decrease in protein levels, p=0.02) in the HOTAIR-tethered cells lacking the essential PRC2 subunit EED (Figure 7D). Knockdown of YTHDC1 resulted in significantly higher luciferase RNA levels in these cells (~2.1 fold change, p=2.2e-05) (Figure 7E). Luciferase enzymatic activity also increased upon YTHDC1 knockdown (~1.3 fold change, p=0.03) (Figure 7F). YTHDC1 knockdown did not affect *HOTAIR* RNA levels in this context (~1.2-fold increase, p=0.8) (Figure 7 – figure supplement 1A-B), indicating that the effects observed on luciferase expression were likely due to disruption of the HOTAIR gene repression mechanism via depletion of YTHDC1 protein rather than loss of HOTAIR expression.

**Figure 7.**
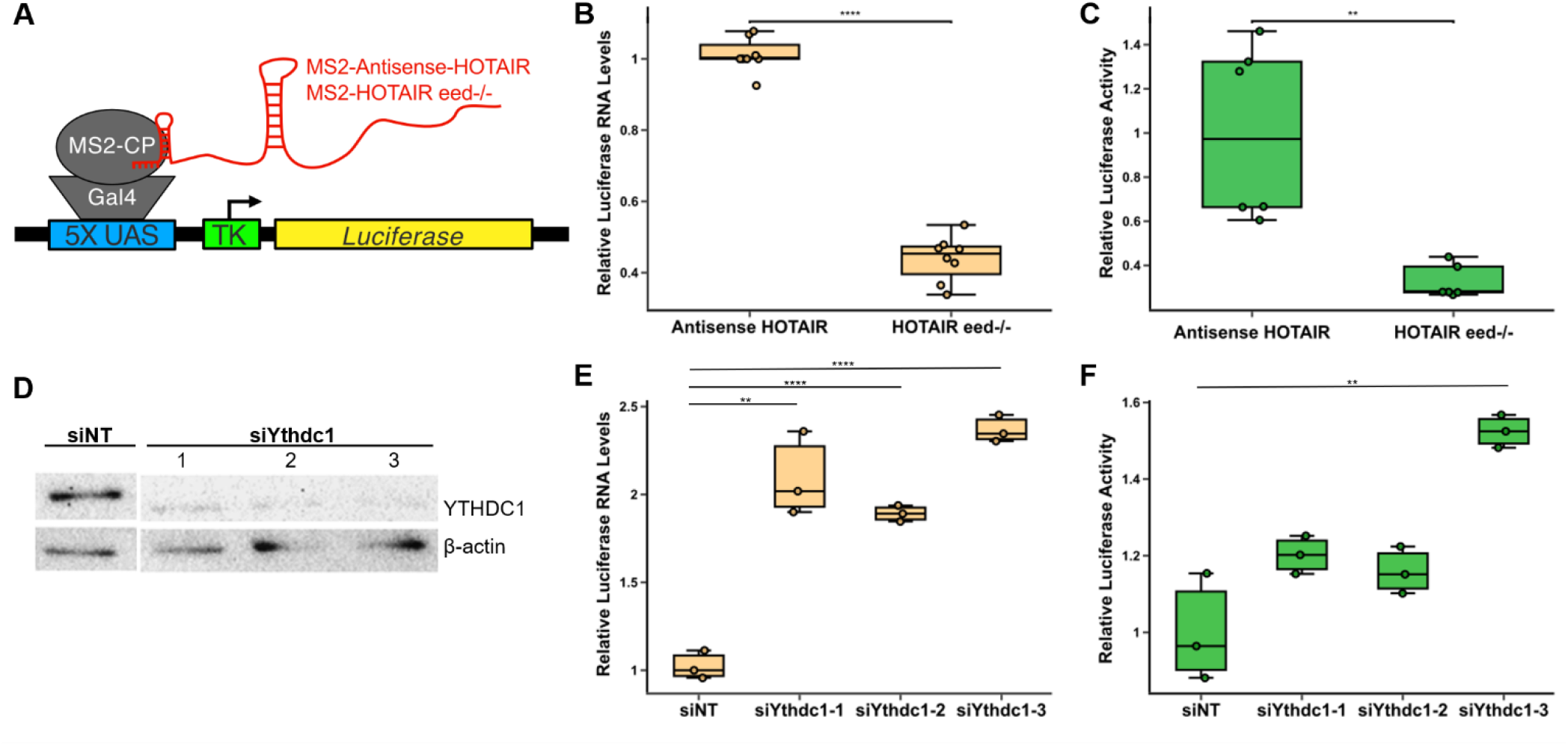
YTHDC1 mediates transcriptional repression by HOTAIR. A) Schematic of 293T cells containing MS2-Antisense-HOTAIR or MS2-HOTAIR tethered upstream of a luciferase reporter. MS2-HOTAIR tethered cells also contain a deletion of *EED*, a subunit of PRC2 that is critical for H3K27 methylation. B-C) Relative luciferase RNA levels (B) and relative luciferase activity (C) in MS2-Antisense HOTAIR or MS2-HOTAIR eed−/− cell lines. D) Western blot of YTHDC1 in MS3-HOTAIR eed−/− 293T reporter cells transfected with non-targeting siRNA or 3 different siRNAs targeting YTHDC1. E-F) Relative luciferase RNA levels (E) and relative luciferase activity (F) of HOTAIR-tethered eed−/− cells transfected with a non-targeting siRNA or 3 different siRNAs against YTHDC1.

## Discussion

Similar to m6A regulation of mRNAs, it is becoming evident that m6A on lncRNAs is both functionally diverse and context dependent. Here, we demonstrate that m6A and the m6A reader YTHDC1 function to enable transcriptional repression by HOTAIR which is analogous to one of the repressive functions demonstrated for the lncRNA Xist(Nesterova et al., 2019; Patil et al., 2016). Our results reveal a mechanism whereby m6A modification of HOTAIR at a specific adenosine residue mediates interaction with YTHDC1, in turn enabling transcriptional interference by HOTAIR which enhances TNBC properties including proliferation and invasion.

### Function of specific m6A sites in HOTAIR

While several m6A sites were identified within HOTAIR when overexpressed, we only detected one m6A site in the endogenously expressed context in MCF-7 cells, making it the most consistently present methylation site. MDA-MB-231 cells overexpressing HOTAIR containing a mutation of this single m6A-modified adenosine had a defect in HOTAIR-mediated proliferation and invasion, as well as its ability to induce HOTAIR-mediated gene expression changes. While A783U mutant HOTAIR appears to retain m6A modification at other sites and interaction with YTHDC1 *in vivo*, we note that the *in vivo* analysis employs formaldehyde crosslinking prior to immunoprecipitation with the YTHDC1 antibody, which enables detection of both weak and strong interactions. It is possible that the A783 m6A site specifically is a high-affinity site for YTHDC1 interaction based on our *in vitro* analysis where YTHDC1 association with methylated domain 2 of HOTAIR was dependent on this site (Figure 5A-C). In line with this hypothesis, we observe a decrease in chromatin-association when A783 of HOTAIR is mutated, which is recovered upon overexpression or direct tethering of YTHDC1 (Figures 5 and 6).

While it is evident that m6A modification of A783 in HOTAIR is important for mediating its effects in breast cancer, other m6A sites within HOTAIR appear to play a role in enabling its high expression levels, potentially through transcript stabilization. When we bypass the normal mechanism of chromatin association using a direct tethering approach for HOTAIR (Figure 7A)(Portoso et al., 2017), YTHDC1 is no longer required for chromatin association or stability, yet is required for gene repression, suggesting a direct role in shutting down transcription, perhaps with LSD1 involvement(Jarroux et al., 2021; Tsai et al., 2010). Our work emphasizes the importance of studying the function of individual m6A sites, as each m6A site has the potential to contribute to the function of an RNA in different ways.

### m6A in gene repression and heterochromatin formation

HOTAIR and other lncRNAs make many dynamic and multivalent interactions with proteins that interact with other proteins, RNA molecules, and chromatin. In the nucleus, the METTL3/14 complex and YTHDC1 are key interactors with m6A-modified RNA that have been shown to regulate chromatin. Work in mouse embryonic stem cells has shown that METTL3 interacts with the SETD1B histone modifying complex, and this plays a role in repression of specific families of endogenous retroviruses(Xu et al., 2021). However, due to the nature of HOTAIR’s mechanism of repressing genes in *trans*, it is unlikely that the METTL3/14 complex remains bound to HOTAIR to induce repression of target loci. For YTHDC1, recent work has found that RNA interactions with this protein can directly regulate chromatin via recruitment of KDM3B, promoting H3K9me2 demethylation and gene expression (Y. Li et al., 2020).In contrast, this study demonstrates that YTHDC1 can act to regulate chromatin association and transcriptional repression by HOTAIR, although the mechanism by which this is accomplished remains ambiguous. Our data suggest that YTHDC1-mediated transcriptional repression occurs upstream of chromatin modification by PRC2. This supports the mechanism of transcriptional interference by HOTAIR proposed by Portoso et. al.(Portoso et al., 2017) (Figure 1A) and suggests that YTHDC1 is an important factor that mediates repression by HOTAIR. Yet, it is still unclear how YTHDC1 binding to a repressive lncRNA mediates transcriptional interference and repression.

### Divergence in m6A and YTHDC1 function for different classes of RNAs

The role of YTHDC1 in mediating chromatin association of and repression by HOTAIR is interesting in the context of the recently identified broad nuclear role of YTHDC1 in regulation of transcription and chromatin state in mouse embryonic stem cells(Jun Liu et al., 2020). While in this case it was demonstrated that YTHDC1 mediates degradation of m6A-modified chromatin-associated regulatory RNAs, our work raises the possibility that YTHDC1 might also mediate transcriptional repression and/or heterochromatin directly through interaction with regulatory RNAs. Our work also shows that, rather than degradation of HOTAIR, m6A sites in HOTAIR mediate its high expression in breast cancer via YTHDC1. An important distinction between HOTAIR and the chromatin-associated regulatory RNAs investigated by Liu *et. al*(Jun Liu et al., 2020) is that HOTAIR regulates genes in *trans* versus *cis*. Liu *et. al* found that YTHDC1 mediates degradation of *cis*-regulatory RNAs by the NEXT complex to slow downstream transcription of their target genes; however, in the case of HOTAIR, YTHDC1 mediates chromatin association of a *trans-*regulatory lncRNA, presumably helping it to repress its target genes in *trans*. We hypothesize that chromatin association of HOTAIR stabilizes it because stable retention of HOTAIR on chromatin as heterochromatin forms is likely to make it inaccessible to factors that mediate its degradation. Our experiments where HOTAIR is tethered to chromatin in a reporter cell line illustrates this, as knockdown of YTHDC1 did not alter the stability of HOTAIR in the context where it is constitutively tethered to chromatin (Figure 6 – figure supplement 1C). It is likely that YTHDC1 performs multiple functions within the nucleus, and that its effects on its target RNAs are context dependent, such as on other nearby RNA binding proteins and/or local chromatin state.

Our work also highlights the fate of HOTAIR-YTHDC1 interaction which is distinctly different from mRNAs whose nuclear export is mediated by YTHDC1(Roundtree et al., 2017). In contrast, we show that YTHDC1 mediates chromatin association of the primarily nuclear-localized HOTAIR lncRNA. Also, while one specific m6A site at A783 is important for mediating chromatin association and the physiological effects of HOTAIR in breast cancer, other m6A sites play a role in overall expression or stability (Figure 8). It is likely that the RNA context and other proteins that either interact directly with YTHDC1 or the RNA molecules it binds to dictate the effects of YTHDC1 binding to its targets. Another possibility is that specific modifications on YTHDC1, such as phosphorylation (which has previously been demonstrated to regulate its localization)(Rafalska et al., 2004), result in interactions with different types of regulatory RNAs or even different m6A sites within a single RNA, ultimately leading to differing effects (i.e. stability vs. chromatin association vs. degradation). Additional studies on how YTHDC1 interacts with specific RNA targets, chromatin, and other proteins in the nucleus will shed light on the mechanisms of YTHDC1 in chromatin regulation.

**Figure 8.**
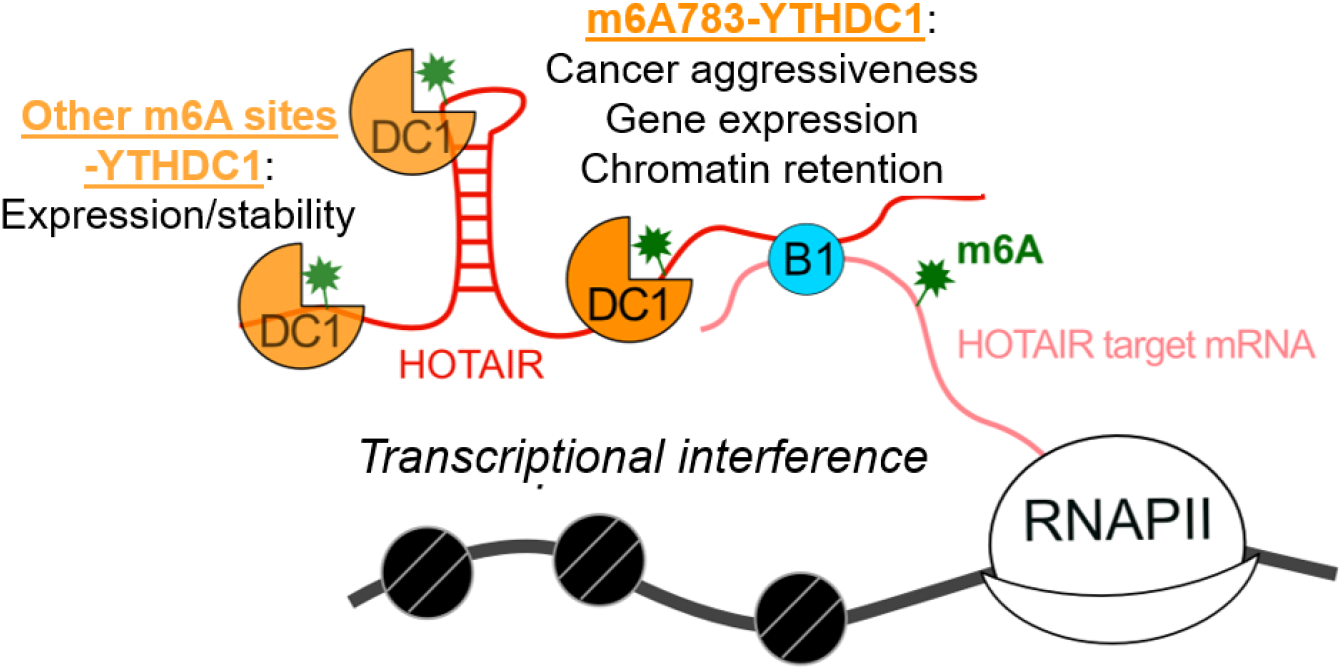
Model of m6A and YTHDC1 effects on HOTAIR. The main function of YTHDC1 occurs via interaction with m6A783 in HOTAIR and mediates chromatin association of HOTAIR to induce transcriptional interference of its target genes, promoting breast cancer growth. YTHDC1 also interacts with other m6A sites within HOTAIR and may mediate its high expression levels and/or stability.

### Antimorphic transformation of HOTAIR function via mutation of a single m6A site

The antimorphic effect of mutating A783 in HOTAIR induced opposite and additional gene expression changes that ultimately resulted in a less aggressive breast cancer state (Figures 1 and 2). Our results show that disruption of a single m6A site can convert HOTAIR from eliciting pro- to anti-tumor effects, allowing overexpression of the converted lncRNA to decrease cancer phenotypes more so than depletion of the wild-type version. Understanding the mechanism behind this induction of antimorphic behavior by a single nucleotide mutation and its biological implications will require future work. Altogether, these findings suggest a potential therapeutic approach to oncogenic lncRNAs such as HOTAIR, where disruption of RNA methylation alone has a greater impact than simple elimination of the RNA.

## Conclusion

The context dependency of m6A function is an emerging theme. With various roles in pluripotency and development and in disease states such as cancer, m6A on different RNA molecules regulates their fate and functional output in different ways(Meyer et al., 2012; L. Wu, Wu, Ning, Liu, & Zhang, 2019). Our work illustrates the context of three specific m6A functions: enabling chromatin association, promoting high levels of lncRNA expression, and facilitating transcriptional repression. We further highlight the importance of one specific m6A site within a lncRNA that contains multiple sites of modification.

As the only primarily nuclear m6A reader, YTHDC1 has the potential to interact with m6A-containing RNA molecules at the site of transcription on chromatin. The outcome of this interaction appears to be dictated by the identity of the RNA molecule (including whether it functions in *cis* or *trans*) and the cell type that it occurs in, but ultimately has the potential to result in chromatin regulation. Our work demonstrates the importance of YTHDC1 in mediating HOTAIR chromatin association and transcriptional repression independent of PRC2, revealing a new layer of regulation by m6A at a specific residue within HOTAIR. Overall, this provides insight into mechanisms of how m6A regulates HOTAIR-mediated breast cancer metastasis which could ultimately lead to new treatment options (for example, preventing m6A methylation at this specific site) for patients with tumors that have elevated HOTAIR levels.

## Materials and Methods

### Cell Culture

MCF-7 cells were maintained in RPMI media (11875093, Invitrogen) and MDA-MB-231 and 293T in DMEM media (MT10013CV, Fisher Scientific). Media contained 10% FBS (F2442-500ML, Sigma-Aldrich) and Pen-Strep (MT30002CI, Fisher Scientific) and cells were grown under standard tissue culture conditions. Cells were split using Trypsin (MT25053CI, Fisher Scientific) according to manufacturer’s instructions.

MDA-MB-231 cells overexpressing WT HOTAIR, A783U mutant HOTAIR, or Anti-Luciferase were generated as previously described using retroviral transduction(Meredith et al., 2016). Stable knockdown of METTL3, METTL14, WTAP, and YTHDC1 and overexpression of YTHDC1 was performed by lentivirus infection of MCF-7 or MDA-MB-231 cells overexpressing HOTAIR or A783U mutant HOTAIR via Fugene HD R.8 with pLKO.1-blasticidin shRNA constructs or a pLX304 overexpression construct as noted in Table S3. Cells were selected with 5 μg/mL blasticidin (Life Technologies). The nontargeting shRNA pLKO.1-blast-SCRAMBLE was obtained from Addgene (Catalog #26701). Two shRNAs for each target were obtained and stable lentiviral transductions with the targeted shRNAs and the scramble control were performed. Cell lines with the most efficient knockdown as determined by western blot were selected for downstream experiments.

### Plasmid Construction

The pBABE-puro retroviral vector was used for overexpression of lncRNAs. The spliced HOTAIR transcript (NR_003716.3) was synthesized and cloned into the pBABE-puro retroviral vector by GenScript. An antisense transcript of the firefly luciferase gene (AntiLuc) was amplified from the pTRE3G-Luciferase plasmid (Clonetech), then cloned into the pBABE-puro retroviral vector. These were generated in a previous publication(Meredith et al., 2016).

To create the A783U mutant HOTAIR overexpression plasmid, staggered QuikChange oligos AG66/AG67 were used to generate the A783U mutation in pTRE3G-HOTAIR using the QuikChange Site Directed Mutagenesis Kit (Agilent 200519) to generate pTRE3G-A783U_HOTAIR. A 1.6Kb fragment of A783U mutant HOTAIR was amplified with primers AG68/AG69 from pTRE3G-A783U_HOTAIR for cloning into pBABE-Puro-HOTAIR cut with XcmI and BamHI by Gibson Assembly. Oligonucleotide sequences are noted in Table S4. All constructs were confirmed by sequencing. pBABE-Puro-6xAU_HOTAIR and pBabe-Puro-14xAU_HOTAIR were synthesized and cloned by GenScript.

Plasmids for the knockdown of METTL3, METTL14, WTAP, and YTHDC1 were generated by cloning the shRNA (RNAi Consortium shRNA Library) from pLKO.1-puro into the pLKO.1-blast backbone (Addgene #26655).

To generate the plasmid for tethering YTHDC1 to HOTAIR via dCasRX, we first constructed a pCDNA-FLAG plasmid by inserting a 5xFLAG sequence (synthesized as a gBlock by IDT DNA) into the HindIII/XbaI site of pCDNA3 (Invitrogen). YTHDC1 was then amplified from pLX304-YTHDC1 (ORF clone ccsdBroad304_04559) with oligonucleotides noted in Table S4, and cloned into the KpnI/NotI site of pCDNA-FLAG to generate pCDNA-FLAG-YTHDC1 (pAJ367). The FLAG-YTHDC1 sequence was amplified then cloned downstream of dCasRX at NheI in the pXR002 plasmid (pXR002: EF1a-dCasRx-2A-EGFP was a gift from Patrick Hsu (Addgene plasmid # 109050; http://n2t.net/addgene:109050; RRID:Addgene_109050)) using oligonucleotides noted in Table S4. Expression of the dCasRX-YTHDC1 fusion protein was confirmed by transfection of the plasmid followed by Western Blot with anti-FLAG M2 mouse monoclonal antibody (F1804, Sigma-Aldrich) and anti-YTHDC1 (14392-1-AP, Proteintech). Plasmids containing guide RNAs were generated using the pXR003 backbone plasmid (pXR003: CasRx gRNA cloning backbone was a gift from Patrick Hsu (Addgene plasmid # 109053; http://n2t.net/addgene:109053; RRID:Addgene_109053)) cut with BbsI, using oligonucleotides noted in Table S4. All plasmids were confirmed by sequencing.

### m6A enhanced crosslinking immunoprecipitation

#### polyA isolation and RNA fragmentation

For each experiment, approximately 100 μg of total RNA was isolated from cells with TRIzol according to manufacturer’s instructions. 10 μg PolyA RNA was isolated using Magnosphere® Ultrapure mRNA Purification Kit (Takara) according to manufacturer’s instructions. PolyA RNA was ethanol precipitated with 2.5 M Ammonium Acetate and 70% ethanol in a solution containing 50 μg/ml GlycoBlue Co-precipitant (AM9515, Invitrogen). RNA was resuspended in 10 μl and fragmented with 10x Fragmentation Buffer (AM8740, Invitrogen) at 75°C for 8 minutes and immediately quenched with 10x Stop Reagent (AM8740, Invitrogen) and placed on ice to generate fragments 30-150 nucleotides in length.

#### Anti-m6A-RNA crosslinking and bead conjugation

Crosslinked RNA-Antibody was generated as previously described(Grozhik, Linder, Olarerin-George, & Jaffrey, 2017). Fragmented RNA was resuspended in 500 μl Binding/Low Salt Buffer (50 mM Tris-HCl pH 7.4, 150 mM Sodium Chloride, 0.5% NP-40) containing 2 μl RNase Inhibitor (M0314, NEB) and 10 μl m6A antibody (ab151230, Abcam), and incubated for 2 hours at room temperature with rotation. RNA-Antibody sample was transferred to one well of a 12-well dish and placed in a shallow dish of ice. Sample was crosslinked twice at 150 mJ/cm^2^ using a Stratagene Stratalinker UV Crosslinker 1800 and transferred to a new tube. 50 μl Protein A/G Magnetic Beads (88803, Pierce) were washed twice with Binding/Low Salt Buffer, resuspended in 100 μl Binding/Low Salt Buffer, and added to crosslinked RNA-Antibody sample. Beads were incubated at 4°C overnight with rotation.

#### eCLIP library preparation

RNA was isolated and sequencing libraries were prepared using a modified enhanced CLIP protocol(Van Nostrand et al., 2016). Beads were washed twice with High Salt Wash Buffer (50 mM Tris-HCl pH 7.4, 1 M Sodium Chloride, 1 mM EDTA, 1% NP-40, 0.5% Sodium Deoxycholate, 0.1% Sodium Dodecyl Sulfate), once with Wash Buffer (20 mM Tris-HCl pH 7.4, 10 mM Magnesium Chloride, 0.2% Tween-20), once with Wash Buffer and 1x Fast AP Buffer (10 mM Tris pH 7.5, 5 mM Magnesium Chloride, 100 mM Potassium Chloride, 0.02% Triton X-100) combined in equal volumes, and once with 1x Fast AP Buffer. Beads were resuspended in Fast AP Master Mix (1x Fast AP Buffer containing 80U RNase Inhibitor (M0314, NEB), 2U TURBO DNase (AM2238, Invitrogen), and 8U Fast AP Enzyme (EF0654, Thermo Scientific)) was added. Samples were incubated at 37°C for 15 minutes shaking at 1200 rpm. PNK Master Mix (1x PNK Buffer (70 mM Tris-HCl pH 6.5, 10 mM Magnesium Chloride), 1 mM Dithiothreitol, 200U RNase Inhibitor, 2U TURBO DNase, 70U T4 PNK (EK0031, Thermo Scientific)) was added to the samples and they incubated at 37°C for 20 minutes shaking at 1200 rpm.

Beads were washed once with Wash Buffer, twice with Wash Buffer and High Salt Wash Buffer mixed in equal volumes, once with Wash Buffer, once with Wash Buffer and 1x Ligase Buffer (50 mM Tris pH 7.5, 10 mM Magnesium Chloride) mixed in equal volumes, and twice with 1x Ligase Buffer. Beads were resuspended in Ligase Master Mix (1x Ligase Buffer, 1 mM ATP, 3.2% DMSO, 18% PEG 8000, 16U RNase Inhibitor, 75U T4 RNA Ligase I (M0437, NEB)), two barcoded adaptors were added (X1a and X1b, see Table S5), and samples were incubated at room temperature for 75 minutes with flicking every 10 minutes. Beads were washed once with Wash Buffer, once with equal volumes of Wash Buffer and High Salt Wash Buffer, once with High Salt Wash Buffer, once with equal volumes of High Salt Wash Buffer and Wash Buffer, and once with Wash Buffer. Beads were resuspended in Wash Buffer containing 1x NuPAGE LDS Sample Buffer (NP0007, Invitrogen) and 0.1M DTT, and incubated at 70°C for 10 minutes shaking at 1200 rpm.

Samples were cooled to room temperature and supernatant was ran on Novex NuPAGE 4-12% Bis-Tris Gel (NP0321, Invitrogen). Samples were transferred to nitrocellulose membrane, and membranes were cut and sliced into small pieces between 20 kDa and 175 kDa to isolate RNA-antibody complexes. Membrane slices were incubated in 20% Proteinase K (03508838103, Roche) in PK Buffer (100 mM Tris-HCl pH 7.4, 50 mM NaCl, 10 mM EDTA) at 37°C for 20 minutes shaking at 1200 rpm. PK Buffer containing 7M urea was added to samples and samples were incubated at 37°C for 20 minutes shaking at 1200 rpm. Phenol:Chloroform:Isoamyl Alcohol (25:24:1) (P2069, Sigma-Aldrich) was added to samples and samples were incubated at 37°C for 5 minutes shaking at 1100 rpm. Samples were centrifuged 3 minutes at 16,000 x *g* and aqueous layer was transferred to a new tube.

RNA was isolated using RNA Clean & Concentrator-5 Kit (R1016, Zymo) according to manufacturer’s instructions. Reverse transcription was performed using AR17 primer (Table S5) and SuperScript IV Reverse Transcriptase (18090010, Invitrogen). cDNA was treated with ExoSAP-IT Reagent (78201, Applied Biosystems) at 37°C for 15 minutes, followed by incubation with 20 mM EDTA and 0.1M Sodium Hydroxide at 70°C for 12 minutes. Reaction was quenched with 0.1M Hydrochloric Acid. cDNA was isolated using Dynabeads MyONE Silane (37002D, ThermoFisher Scientific) according to manufacturer’s instructions. 20% DMSO and rand3Tr3 adaptor (Table S5) was added to samples, and samples were incubated at 75° for 2 minutes. Samples were placed on ice and Ligation Master Mix (1x NEB Ligase Buffer, 1mM ATP, 25% PEG 8000, 15U T4 RNA Ligase I (NEB)) was added to samples. Samples were mixed at 1200 rpm for 30 seconds prior to incubation at room temperature overnight.

cDNA was isolated using Dynabeads MyONE Silane according to manufacturer’s instructions and eluted with 10 mM Tris-HCl pH 7.5. A 1:10 dilution of cDNA was used to quantify the cDNA library by qPCR using a set of Illumina’s HT Seq primers, and Ct values were used to determine number of cycles for PCR amplification of cDNA. The undiluted cDNA library was amplified by combining 12.5μL of the sample with 25μL Q5 Hot Start PCR Master Mix and 2.5μL (20μM) of the same indexed primers used previously. Amplification for the full undiluted sample used 3 cycles less than the cycle selected from the diluted sample. The PCR reaction was isolated using HighPrep PCR Clean-up System (AC-60050, MAGBIO) according to manufacturer’s instructions.

The final sequencing library was gel purified by diluting the sample with 1x Orange G DNA loading buffer and running on a 3% quick dissolve agarose gel containing SYBR Safe Dye (1:10,000). Following gel electrophoresis, a long wave UV lamp was used to extract DNA fragments from the gel ranging from 175 to 300 base pairs. The DNA was isolated using QiaQuick MinElute Gel Extraction Kit (28604, Qiagen). The purified sequencing library was analyzed via TapeStation using DNA ScreenTape (either D1000 or HS D1000) according to the manufacturer’s instructions to assess for appropriate size and concentration (the final library should be between 175 and 300 base pairs with an ideal concentration of at least 10nM).

#### Sequencing and analysis

Samples were sequenced at the Genomics and Microarray Shared Resource facility at University of Colorado Denver Cancer Center on an Illumina MiSeq or NovaSEQ6000 with 2x 150 base pair paired-end reads to generate 40 million raw reads for each sample. Computational analysis methods are described in (Roberts et al., 2020). Briefly, a custom Snakemake workflow was generated based on the original eCLIP analysis strategies(Van Nostrand et al., 2016) to map reads to the human genome. To identify m6A sites, we used a custom analysis pipeline to identify variations from the reference genome at single-nucleotide resolution across the entire genome. We then employed an internally developed Java package to identify C-to-T mutations occurring 1) within the m^6^A consensus motif ‘RAC’: ‘R’ is any purine, A or G; A being the methylated adenosine; and C where the mutation occurs; and 2) within a frequency range of greater than or equal to 2.5% and less than or equal to 50% of the total reads at a given position (with a minimum of 3 C-to-T mutations at a single site). The resulting m^6^A sites were then compared to those identified in the corresponding input sample and any sites occurring in both were removed from the final list of m^6^A sites (this eliminates any mutations that are not directly induced from the anti-m^6^A antibody crosslinking). Full transcriptome data associated with the methods manuscript(Roberts et al., 2020) is at GEO accession number GSE147440.

### m6A RNA Immunoprecipitation (meRIP)

Total RNA was isolated with TRIzol (15596018, Invitrogen) according to the manufacturer’s instructions. RNA was diluted to 1 μg/μl and fragmented with 1x Fragmentation Buffer (AM8740, Invitrogen) at 75°C for 5 minutes. 1x Stop Reagent (AM8740, Invitrogen) was added immediately following fragmentation and samples placed on ice. 500 ng of input sample was reserved in 10 μl nuclease free water for qRT-PCR normalization. Protein A/G Magnetic Beads (88803, Pierce) were washed twice with IP Buffer (20 mM Tris pH 7.5, 140 mM NaCl, 1% NP-40, 2mM EDTA) and coupled with anti-m6A antibody (ab151230, Abcam) or an IgG control (NB810-56910, Novus) for 1 hour at room temperature. Beads were washed 3 times with IP Buffer. 10 μg fragmented RNA and 400U RNase inhibitor was added to 1 ml IP Buffer. Antibody-coupled beads were resuspended in 500 μl RNA mixture and incubated 2 hours to overnight at 4°C on a rotor. Beads were washed 5 times with cold IP Buffer. Elution Buffer (1x IP Buffer containing 10 U/μl RNase inhibitor and 0.5 mg/ml N6-methyladenosine 5’-monophosphate (M2780, Sigma-Aldrich) was prepared fresh and kept on ice. Samples were eluted with 200 μl Elution Buffer for 2 hours at 4°C on a rotor. Supernatant was removed and ethanol precipitated with 2.5M Ammonium Acetate, 70% Ethanol, and 50 μg/ml GlycoBlue Coprecipitant (Invitrogen AM9515). RNA was washed with 70% ethanol, dried for 10 minutes at room temperature, and resuspended in 10 μl nuclease free water. RNA was quantified by nanodrop and 200 ng RNA was reverse transcribed using High Capacity cDNA Reverse Transcription Kit (4368814, ThermoFisher Scientific) and quantified by qPCR (oligonucleotides listed in Table S6), and fraction recovered was calculated from Input and IP values.

### RNA Immunoprecipitation of YTHDC1

Actively growing cells from 70-90% confluent 15-cm dishes were trypsinized and washed twice with ice-cold 1x PBS. Cell pellet was resuspended in 1% V/V Formaldehyde (28908, Pierce) in 1x PBS and incubated at room temperature for 10 minutes on a rotor. Crosslinking was quenched with 0.25 M glycine at room temperature for 5 minutes. Cells were washed 3 times with ice-cold 1x PBS and placed on ice. 20 μl Protein A/G beads were washed twice with RIPA Binding Buffer (50 mM Tris-HCl pH 7.4, 100 mM Sodium Chloride, 1% NP-40, 0.1% Sodium Dodecyl Sulfate, 0.5% Sodium Deoxycholate, 4 mM Dithiothreitol, 1x Protease Inhibitors), resuspended in 1 ml RIPA Binding Buffer, and split to two 0.5 ml aliquots. 2 μg YTHDC1 antibody (ab122340, Abcam) or an IgG Control (sc-2027, Santa Cruz Biotechnology) was added to beads and incubated for 2 hours at 4°C on a rotor. Fixed cells were resuspended in 1 ml RIPA Binding Buffer and placed in the Bioruptor Pico (B01060010, Diagenode) for 10 cycles of 30 seconds on, 30 seconds off. Lysates were digested with TURBO DNase for 5 minutes at 37°C with mixing at 1000 rpm and transferred to ice for 5 minutes. Lysates were clarified by centrifugation at 17,000*g* at 4°C for 10 minutes and supernatant was transferred to a new tube. 200U RNase Inhibitor was added to the 1 ml clarified lysate. A 5% aliquot was removed and processed downstream with IP samples. A 2% aliquot was removed and diluted with 1x SDS Sample Buffer (62.5 mM Tris-HCl pH 6.8, 2.5% SDS, 0.002% Bromophenol Blue, 5% β-mercaptoethanol, 10% glycerol) and protein input and recovery was monitored by Western Blot. Antibody-coupled beads were washed 3 times with RIPA Binding Buffer and resuspended in half of the remaining lysate. Samples were incubated overnight at 4°C on a rotor. Beads were washed 5 times with RIPA Wash Buffer (50 mM Tris-HCl pH 7.4, 1 M Sodium Chloride, 1% NP-40, 0.1% Sodium Dodecyl Sulfate, 0.5% Sodium Deoxycholate, 1 M Urea, 1x Protease Inhibitors) and resuspended in 100 μl RNA Elution Buffer (50 mM Tris-HCl pH 7.4, 5 mM EDTA, 10 mM Dithiothreitol, 1% Sodium Dodecyl Sulfate). Input sample was diluted with 1x RNA Elution Buffer. Formaldehyde crosslinks in both input and IP samples were reversed by incubation at 70°C for 30 minutes at 1000 rpm. Supernatant was transferred to a new tube and RNA was isolated using TRIzol-LS according to the manufacturer’s instructions. Reverse transcription was performed on 100 ng RNA using SuperScript IV Reverse Transcriptase. qPCR was performed as described below.

### RNA Isolation and qRT-PCR

RNA was isolated with TRIzol (Life Technologies) with extraction in chloroform followed by purification with the RNeasy kit (Qiagen). Samples were DNase treated using TURBO DNase (Ambion). Reverse transcription was performed using the cDNA High Capacity Kit (Life Technologies). qPCR was performed using Sybr Green master mix (Takyon, AnaSpec Inc.) using the primers listed in Table S6 on a C1000 Touch Thermocycler (BioRad). EEF1A1 primer sequences were obtained from the Magna MeRIP m6A kit (17-10499, Sigma-Aldrich). Sequences for Luciferase primers (LucR2) were obtained from a previous publication(Vaquero et al., 2004). Three qPCR replicates were performed for each sample, and these technical replicates were averaged prior to analysis of biological replicates. At least 3 biological replicates were performed for each qPCR experiment.

### Cell Proliferation Assays

Three independent clones, here defined as a pool of selected cells stably expressing the pBabe plasmid, were analyzed for cell proliferation. 2,000 cells were plated in a 96-well dish in DMEM media containing 10% FBS and selective antibiotics (1μg/ml puromycin (P8833, Sigma-Aldrich) and/or 5μg/ml blasticidin (71002-676, VWR)), allowed to settle at room temperature for 20 minutes, then placed in an Incucyte® S3 (Sartorius). Pictures were taken with a 10x magnification every 2 hours for 48 hours using a Standard scan. Confluency was determined using the Incucyte ZOOM software. Growth rate was calculated from % confluency using the Least Squares Fitting Method(Roth, 2006).

### Cell Invasion Assays

MDA-MB-231 cell lines were grown to 70-90% confluence and serum starved in OptiMEM for ~20 hours prior to setting up the experiment. Cells were washed, trypsinized, and resuspended in 0.5% serum DMEM. 10% serum DMEM was added to the bottom chamber of Corning Matrigel™ Invasion Chambers (Corning 354481), and 200,000 cells were plated in the top chamber in 0.5% serum DMEM. Cells were incubated for 22 hours at 37°C followed by 4% PFA fixation and 0.1% Crystal Violet staining. Matrigel inserts were allowed to dry overnight, followed by brightfield imaging with a 20X air objective. Four biological replicates were performed, with technical duplicates in each set. For each Matrigel insert, four fields of view were captured, and cells were counted in Fiji (eight data points per condition, per biological replicate). The violin plot includes all of the data points, while statistical analysis was performed on the average number of cells/field for each biological replicate.

### Gene Expression Analyses

Total RNA was extracted from MDA-MB-231 cells using TRIZol (Life Technologies) with extraction in chloroform followed by purification with the RNeasy kit (Qiagen). Samples were DNase treated using TURBO DNase (Ambion). polyA-selected sequencing libraries were prepared and sequenced by The Genomics Shared Resource at the University of Colorado Cancer Center. All gene expression data associated with this publication is available through GEO accession number GSE173530. Differential gene expression analysis was performed using Salmon and DESeq2(Love, Huber, & Anders, 2014; Patro, Duggal, Love, Irizarry, & Kingsford, 2017). Briefly, the reads were quantified using salmon to generate transcript abundance estimates and then DESeq2 was used to determine differential expression between samples. Heat maps were generated by using normalized read counts of genes that were significantly (p<0.1) differentially expressed between conditions to generate Z-scores. GO term enrichment analysis was performed using the GO Consortium’s online PANTHER tool(Ashburner et al., 2000; Mi, Muruganujan, Ebert, Huang, & Thomas, 2019; “The Gene Ontology resource: enriching a GOld mine.,” 2021). To analyze correlation between expression in HOTAIR v. Control and A783U v. Control pairwise comparisons, the total set of differentially expressed genes were filtered to include only those whose fold change value was greater than 1.15 in either direction for both comparisons. These values were then plotted against each other. Linear regression was used to fit a trend line over the points, with the calculated Pearson correlation coefficient included in the graph.

### Purification of METTL3/14

Suspension-adapted HEK293 cells (Freestyle™ 293-F cells, R790-07, Life Technologies) were grown as recommended in Freestyle™ 293 Expression Medium (12338026, Life Technologies,) sharking at 37°C in 5% CO_2_. Cells were grown to a concentration of 3 x 10^6^ cells/ml and diluted to 1 x 10^6^ cells/ml in 50 ml 293F Freestyle Media 24 hours prior to transfection. Before transfection, cells were spun down and resuspended in 50 ml fresh 293F Freestyle Media at a concentration of 2.5 x 10^6^ cells/ml. Expression plasmid (pcDNA3.1-FLAG-METTL3, pcDNA3.1-FLAG-METTL14) were added to the flask at a concentration of 1.5 μg, and flask was shaken in the incubator for 5 minutes. 9 μg/ml PEI was added to the flask and cells were returned to incubator. After 24 hours of growth, an additional 50 ml fresh 293F Freestyle Media was added and culture was supplemented with 2.2 mM VPA. Cells were harvested as two 50 ml pellets 72 hours after addition of VPA.

Cell pellets were resuspended in 1x Lysis Buffer (50 mM Tris pH 7.4, 150 mM Sodium Chloride, 1 mM EDTA, 1% TritonX-100, 1x Protease inhibitors) to obtain a concentration of 10^7^ cells/ml and incubated for 20 minutes at 4°C with rotation. Cell lysate was clarified by centrifugation at 4°C, 12,000 x *g* for 15 minutes. Supernatant was transferred to a new tube and kept on ice. Anti-FLAG M2 affinity resin was equilibrated with 1x Lysis Buffer by washing 3 times. Equilibrated resin was resuspended in 1x Lysis Buffer and added to the tube containing the clarified lysate. Sample was incubated for 2 hours at 4°C with rotation. Resin was pelleted by centrifugation at 4°C, 500 x *g*. Supernatant was removed, and resin was washed 3 times with 1x Wash Buffer (50 mM Tris pH 7.4, 150 mM Sodium Chloride, 10% Glycerol, 1 mM Dithiothreitol) for 5 minutes each at 4°C with rotation. Sample was equilibrated to room temperature and resin was resuspended in 1x Wash Buffer containing 0.2 mg/ml 3xFLAG Peptide. Samples were incubated at room temperature for 10 minutes shaking at 1000 rpm, centrifuged for 2 minutes at 1000 x g, and supernatant was reserved (elution 1). Elution was repeated twice to obtain two additional elution samples (elution 2 and 3). Samples were analyzed by Coomassie to determine protein concentration and purity. Samples were aliquoted and stored at −80°C and thawed on ice prior to use in *in vitro* m6A methylation experiments.

### *In vitro* m6A methylation and interaction assays

All plasmids and oligonucleotides used in this assay are listed in Table S7. Using PCR, we generated a DNA fragment for Domain 2 of wild-type (pTRE3G-HOTAIR, pAJ171) and A783U (pTRE3G-A783U_HOTAIR, pAJ385) mutant HOTAIR using primers MB88 and MB89. A 5’ T7 promoter and 3’ RAT tag were added to the sequence via PCR with primers MB22 and MB94. *in vitro* transcription of the PCR templates was completed using the MEGAScript T7 Transcription Kit (AM1334, ThermoFisher Scientific) according to the manufacturer’s instructions, and RNA was purified using the RNeasy Mini Kit (Qiagen 75106). 500 nM RNA was diluted in 1x Methyltransferase Buffer (20 mM Tris pH 7.5, 0.01% Triton-X 100, 1 mM DTT) in reactions containing 50 μM SAM and 500 nM purified METTL3/14 (+m6A) for 1 hour at room temperature. Control reactions contained no METTL3/14 (-m6A). RNA was purified using the RNeasy Mini Kit according to manufacturer’s instructions.

To obtain FLAG-tagged YTHDC1 protein, 293 cells were transfected using Lipofectamine 2000 (11668030, ThermoFisher Scientific) with plasmid YTHDC1-FLAG and cell lysates were generated as previously described(Meredith et al., 2016). Dynabeads (M270, Invitrogen) were resuspended in high-quality dry Dimethylformamide at a concentration of 2 x 10^9^ beads/ml. Dynabeads were stored at 4°C and equilibrated to room temperature prior to use. Dynabeads were washed in 0.1 M Sodium Phosphate Buffer (pH 7.4) and vortexed for 30 seconds. A second wash was repeated with vortexing and incubation at room temperature for 10 minutes with rotation. 1 mg/ml IgG solution was prepared by diluting rabbit IgG (15006, Sigma) in 0.1 M Sodium Phosphate Buffer. Washed beads were resuspended in 0.1 M Sodium Phosphate Buffer at a concentration of 3 x 10^9^ beads/ml, and an equal volume of 1 mg/ml IgG was added. Samples were vortexed briefly and an equal volume of 3M Ammonium Sulfate was added and samples were mixed well. Samples were incubated at 37°C for 18-24 hours with rotation. Samples were washed once briefly with 0.1 M Sodium Phosphate Buffer, then twice with incubation at room temperature for 10 minutes with rotation. Samples were washed in Sodium Phosphate Buffer + 1% TritonX-100 at 37°C for 10 minutes with rotation. A quick wash with 0.1 M Sodium Phosphate Buffer was performed and followed by 4 washes in 0.1 M Citric Acid pH 3.1 at a concentration of 2 x 10^8^ beads/ml at room temperature for 10 minutes with rotation. After a quick wash with 0.1 M Sodium Phosphate Buffer, beads were resuspended to 1 x 10^9^ beads/ml in 1x PBS + 0.02% Sodium Azide and stored at 4°C prior to use.

800 ng of +/−m6A RNA was incubated with 150 ng PrA-PP7 fusion protein in HLB300 (20 mM Hepes pH 7.9, 300 mM sodium chloride, 2 mM magnesium chloride, 0.1% NP-40, 10% glycerol, 0.1 mM PMSF, 0.5 mM DTT). RNA was prebound to PP7 for 30 minutes at 25°C, 1350 rpm. 75 μl IgG-coupled Dynabeads were washed with HLB300 twice and resuspended in 250 μl HLB300. 50 μl beads were added to each tube of RNA-PP7 and samples were incubated 1 hour at 25°C, 1350 rpm. Beads were washed twice with HLB300 and resuspended in 80 μl Binding Buffer (10 mM Hepes pH 7.4, 150 mM potassium chloride, 3 mM magnesium chloride, 2 mM DTT, 0.5% NP-40, 10% glycerol, 1mM PMSF, 1x protease inhibitors) containing 80U RNase Inhibitor. 25 μg YTHDC1-FLAG containing lysate and 800 ng competitor RNA (IVT untagged HOTAIR D2) was added to each sample. Samples were incubated at 4°C for 2.5 hours on a rotor. Beads were washed 3 times with cold Wash Buffer (200 mM Tris-HCl pH 7.4, 200 mM sodium chloride, 2 mM magnesium Chloride, 1 mM DTT, 1x protease inhibitors) and resuspended in 1x SDS loading buffer. A 10% protein input sample was diluted in 1x SDS loading buffer. Samples were boiled 5 minutes at 95°C and supernatant transferred to a new tube. Half of each sample was loaded on a 10% acrylamide gel and Western Blot was performed using anti-FLAG antibody.

### Fractionation

Cells were grown in 15-cm dishes to 70-90% confluency. Cells were released with Trypsin (Corning), washed once with 1x PBS containing 1 mM EDTA, and split into two volumes. 1/4 of the sample was harvested in TRIzol and RNA isolated with RNeasy kit for the input RNA sample. The remaining ¾ of the sample was fractionated into cytoplasmic, nucleoplasmic, and chromatin-associated samples. Cells were lysed in cold Cell Lysis Buffer (10 mM Tris-HCl pH 7.5, 0.15% NP-40, 150 mM Sodium Chloride) containing RNase inhibitors for 5 minutes on ice. Lysate was layered onto 2.5 volumes of Sucrose Cushion (10mM Tris-HCl pH7.5, 150 mM Sodium Chloride, 24% Sucrose) containing RNase inhibitors. Samples were centrifuged for 10 minutes at 17,000*xg* at 4°C. Supernatant was collected (Cytoplasmic sample). Pellet was rinsed with 1x PBS containing 1 mM EDTA and resuspended in cold Glycerol Buffer (20 mM Tris-HCl pH 7.9, 75 mM Sodium Chloride, 0.5 mM EDTA, 0.85 mM DTT, 0.125 mM PMSF, 50% Glycerol) containing RNase inhibitors. An equal volume of cold Nuclei Lysis Buffer (10 mM HEPES pH 7.6, 1 mM DTT, 7.5 mM Magnesium Chloride, 0.2 mM EDTA, 0.3 M Sodium Chloride, 1M Urea, 1% NP-40) was added and sample was briefly vortexed twice for 2 seconds. Samples were incubated on ice 2 minutes and centrifuged for 2 minutes at 17,000*xg* at 4°C. Supernatant was collected (Nucleoplasmic sample). The remaining pellet was resuspended in 1x PBS containing 1 mM EDTA (Chromatin-associated sample). Each sample was subjected to TURBO DNase digestion at 37°C for 30 minutes in 1x TURBO Buffer and 10 U TURBO for cytoplasmic and nucleoplasmic samples, or 40U TURBO for chromatin-associated sample. Reactions were quenched with 10mM EDTA and 3 volumes of TRIzol-LS was added. RNA isolation was performed as recommended by manufacturer. Samples were quantified by nanodrop to determine RNA concentration and ran on a 2% agarose gel to confirm RNA integrity. qRT-PCR was performed on 2 μg of RNA and normalized to RNA recovery, input values, and GAPDH.

### dCasRX-YTHDC1 and gRNA Transfection

One plasmid containing dCasRX-FLAG-YTHDC1 in pXR002 in combination with one plasmid containing the designated guide RNA in pXR003 (see description in Plasmid Construction) were transfected into a 70-90% confluent 10-cm dish using Lipofectamine 2000 (11668030, Invitrogen) according to manufacturer’s instructions. Plates were incubated at 37°C for ~24 hours, then subjected to fractionation as described above.

### siRNA Transfection

Silencer Select siRNAs were obtained from ThermoFisher targeting YTHDC1 (n372360, n372361, n372362) or Negative Controls (4390843, 4390846) and transfected into 293 cell lines using Lipofectamine RNAiMAX Transfection Reagent (13778030, ThermoFisher). Transfections were performed in a 24-well plate with 5 pmol of siRNA and 1.5 μl RNAiMAX Transfection reagent per well. Cells were harvested 24 hours after transfections and analyzed by Luciferase Assay and qRT-PCR.

### Luciferase Assay

Analysis of luciferase activity was performed using the Luciferase Assay System (E1500, Promega). Cells were washed with 1x PBS and lysed in 100 μl 1x Cell Culture Lysis Reagent. Cells were scraped from bottom of dish and suspension was transferred to a new tube. Lysates were frozen and thawed prior to luciferase assay to ensure complete lysis. Luciferase assays were performed on 20 μl of lysate or 1x Cell Culture Lysis Reagent in 96 well plates on the GloMax-Multi Detection System (TM297, Promega). 100 μl Luciferase Assay Reagent was added to wells, mixed, and light production measured. Measurements were performed in 3 technical replicates for each biological replicate. Luciferase activity was normalized to protein concentration of samples.

### Statistical Analyses

Graphs were prepared and data fitting and statistical analyses were performed using Biovinci© (version 1.1.5, Bioturing, Inc., San Diego, California, USA). Each box-and-whisker plot displays datapoints for each replicate, the median value as a line, a box around the lower and upper quartiles, and whiskers extending to maximum and minimum values, excluding outliers as determined by the upper and lower fences. A student’s unpaired T-test was used to determine statistical significance. Differences and relationships were considered statistically significant when *p* ≤ 0.05. For all graphs, * *p* < 0.05, ** *p* < 0.01, *** *p* < 0.001, **** *p* < 0.0001.

## Supporting information

Gene Expression Analysis

## Acknowledgements

This work was funded by the NIH (R35GM119575 (AMJ), R01CA187733 (JKR), T32CA190216 (AMP), T32GM008730 (JTR), F31CA247343 (JTR), F32CA239436 (MMW), and NCI Cancer Center Support Grant P30CA046934 to the University of Colorado Cancer Center), a Department of Defense Breast Cancer Research Program Fellowship Award BC170270 (AMP), and an RNA Bioscience Initiative Grant. We thank Maggie M. Balas for figure advice and preparation, Rafael Margueron for providing MS2-tethered HOTAIR cell lines used in this study, Chuan He for providing METTL3 and METTL14 expression plasmids, Patrick Hsu for providing the dCasRX plasmids, and John Rinn, Suja Jagannathan, Neelanjan Mukherjee, and Maria Aristizabal for their suggestions on the manuscript.

## Author Contributions

AMP, JTR, JKR, and AMJ designed research; AMP, JTR, MC, EDD, AL, MK performed experiments; AMP, JTR, and MMW analyzed data; and AMP, JKR, and AMJ wrote the paper.

---

**Supplemental File 1. Figure supplements.**

**Supplemental File 2. Supplemental tables of m6A sites identified in HOTAIR and ORFs, shRNAs, plasmids, and oligonucleotides used in this study.**

**Supplemental File 3. Excel file of differentially expressed genes identified in DESeq2 pairwise comparisons.**

