## Supplemental Figures for "A single N6-methyladenosine site in lncRNA HOTAIR regulates its function in breast cancer cells"

<u>**Supplemental File 1. Figure Supplements**</u>

**Figure 1 – figure supplement 1**

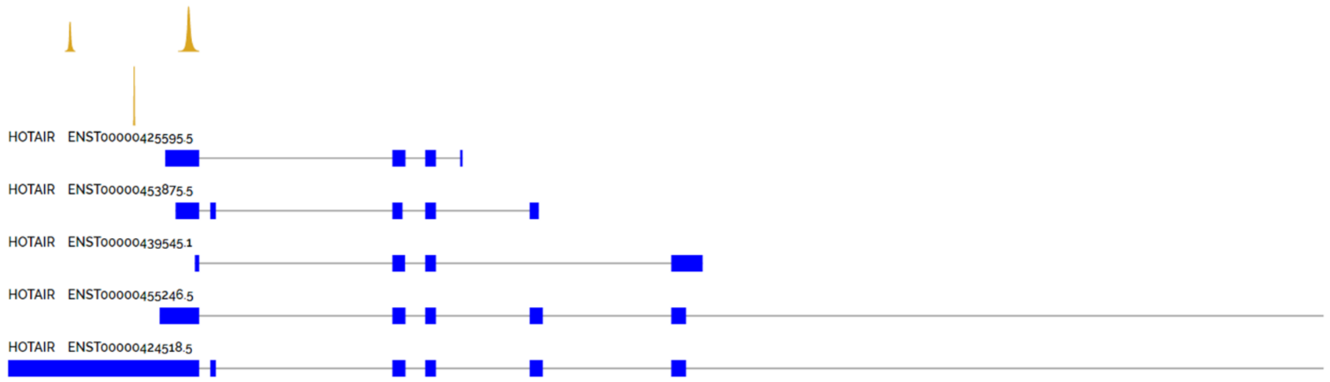


| Peak Region | Strand | Enrichment Score | Gene Symbol | Gene Type | Gene Region | Subcellular Location | Cell Line | Condition | Library Type |
|---|---|---|---|---|---|---|---|---|---|
| chr12:53962847-53962946 | - | 1.5 | HOTAIR | lncRNA | NA | NA | Hela | Cancer | MeRIP-Seq |
| chr12:53963926-53964124 | - | 2.29 | HOTAIR | lncRNA | NA | NA | Hela | Cancer | MeRIP-Seq |
| chr12:53963496-53963516 | - | 3 | HOTAIR | lncRNA | NA | NA | Hela_PA | Cancer | PA-m6A-Seq |

**Figure 1 – figure supplement 1. CVm6A visualization of HOTAIR m6A RIP experiments.** Data obtained from http://gb.whu.edu.cn:8080/CVm6A

### Figure 1 – figure supplement 2

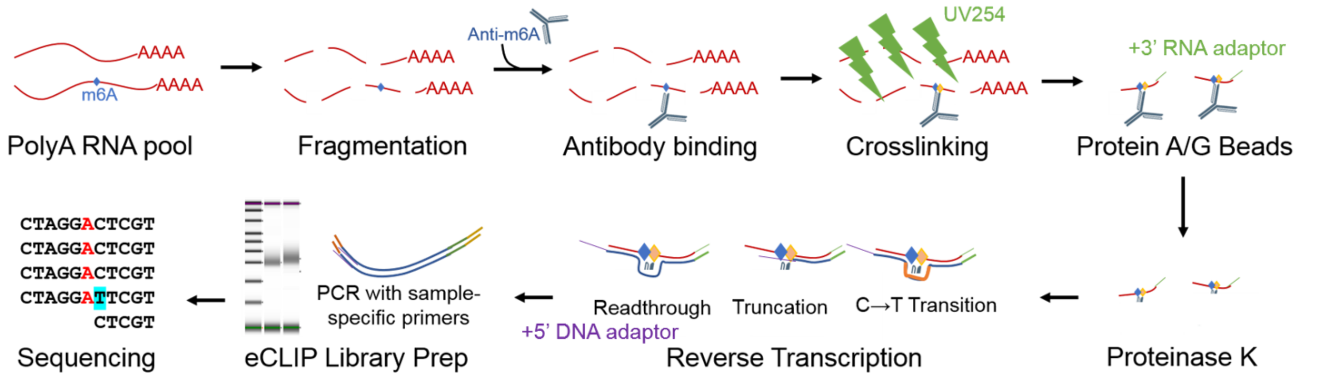


**Figure 1 – figure supplement 2. Strategy for mapping m6A at single nucleotide sites.** Schematic of m6A eCLIP pipeline used to map m6A sites.

### Figure 1 – figure supplement 3

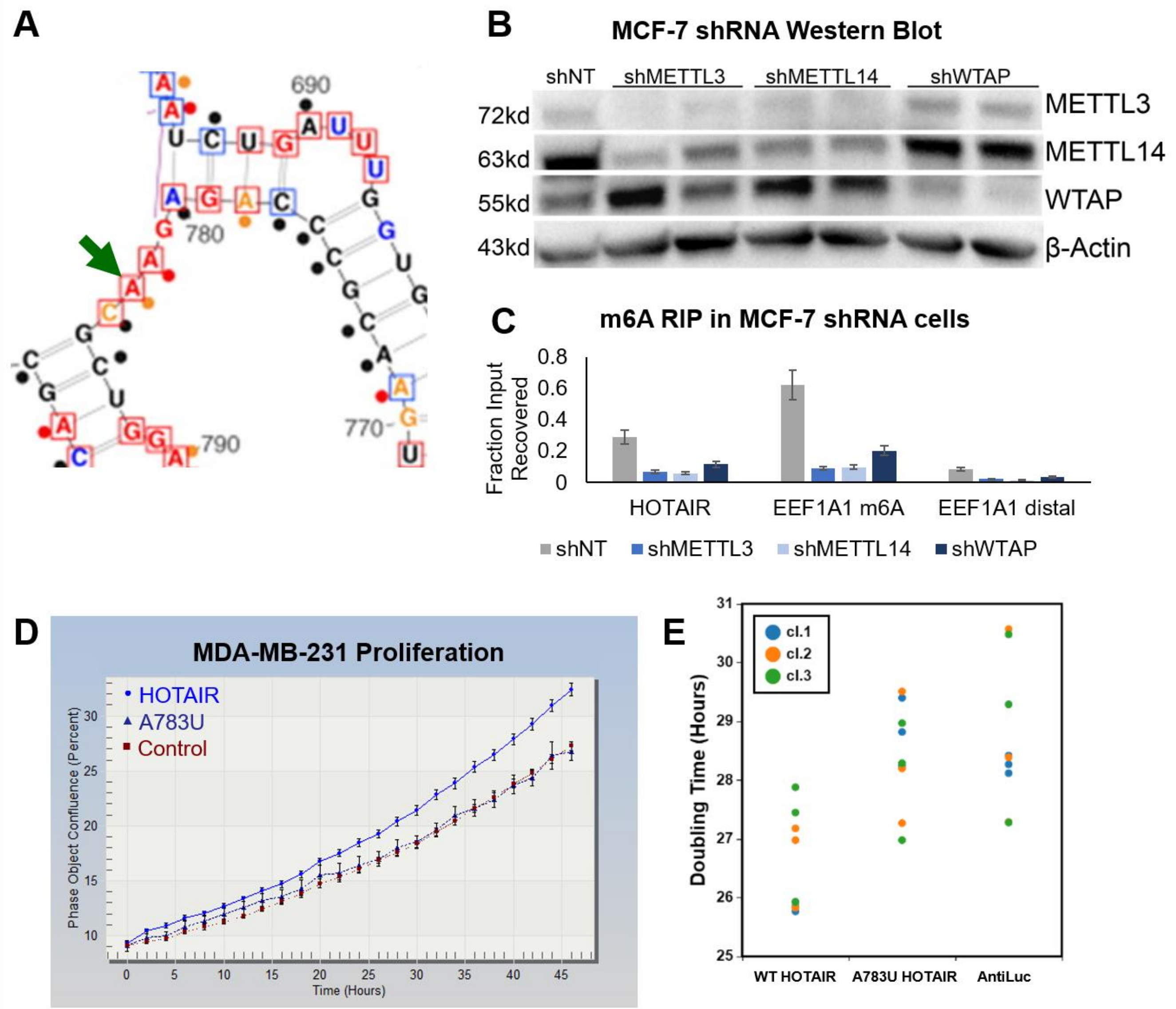


**Figure 1 – figure supplement 3. Nucleotide A783 is m6A modified and important for HOTAIR-mediated breast cancer growth. A)** Portion of HOTAIR structure from (Somarowthu et al., 2015). A783 is marked by a blue arrow. **B)** Western blot of knockdown lines generated in MCF-7 cells. **C)** m6A RIP results on MCF-7 knockdown lines. **D)** Example of growth curve obtained from Incucyte experiments. Percent confluence was measured every 2 hours for 48 hours on MDA-MB-231 cell lines noted. **E)** Data from Figure 1F, doubling time of MDA-MB-231 overexpression cell lines, displayed as individual data points (3 clones for each cell line noted, 3 biological replicates for each clone).

Figure 2 – figure supplement 1

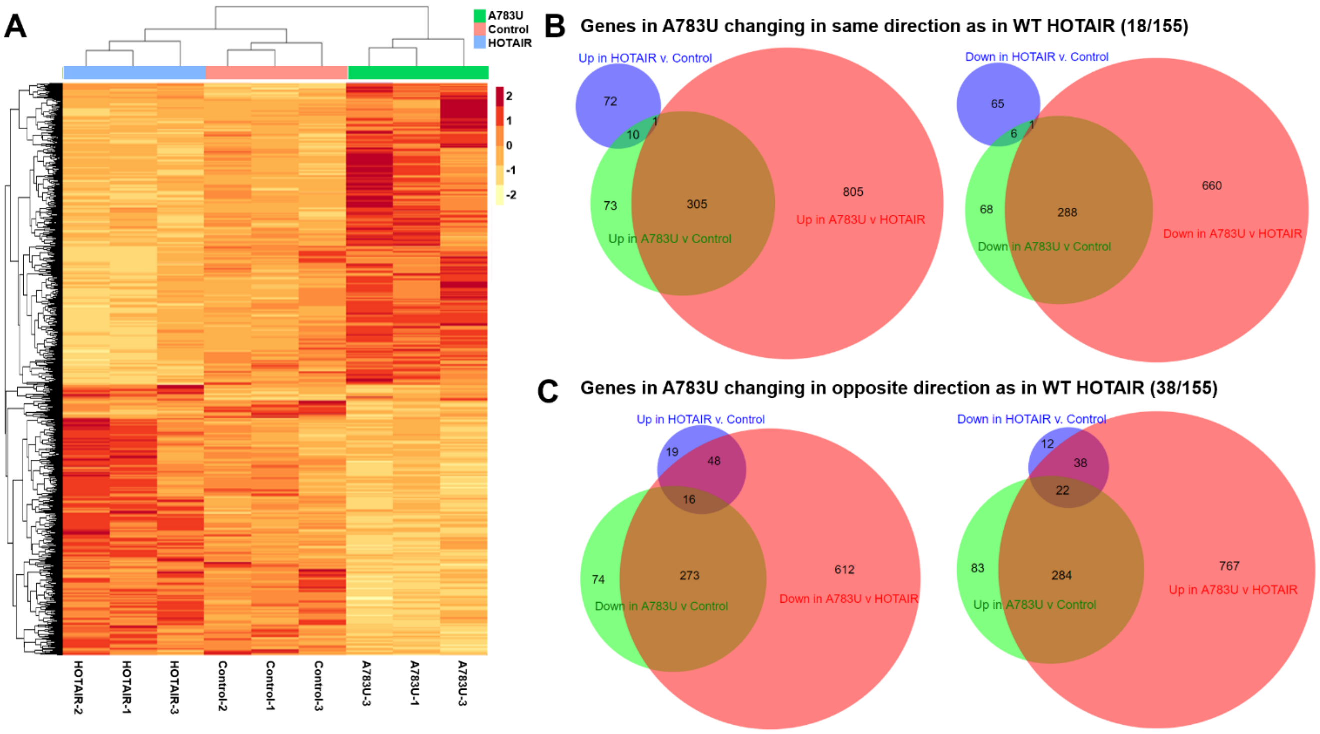


**Figure 2 – figure supplement 1. Expression of A783U HOTAIR induces opposite gene expression changes compared to WT HOTAIR.** (A) Heatmap of Z-scores of all DEGs identified in pairwise comparisons. (B-E) Venn diagrams (created using BioVenn, (Hulsen et al., 2008)) of comparisons of DEGs by expression pattern noted.

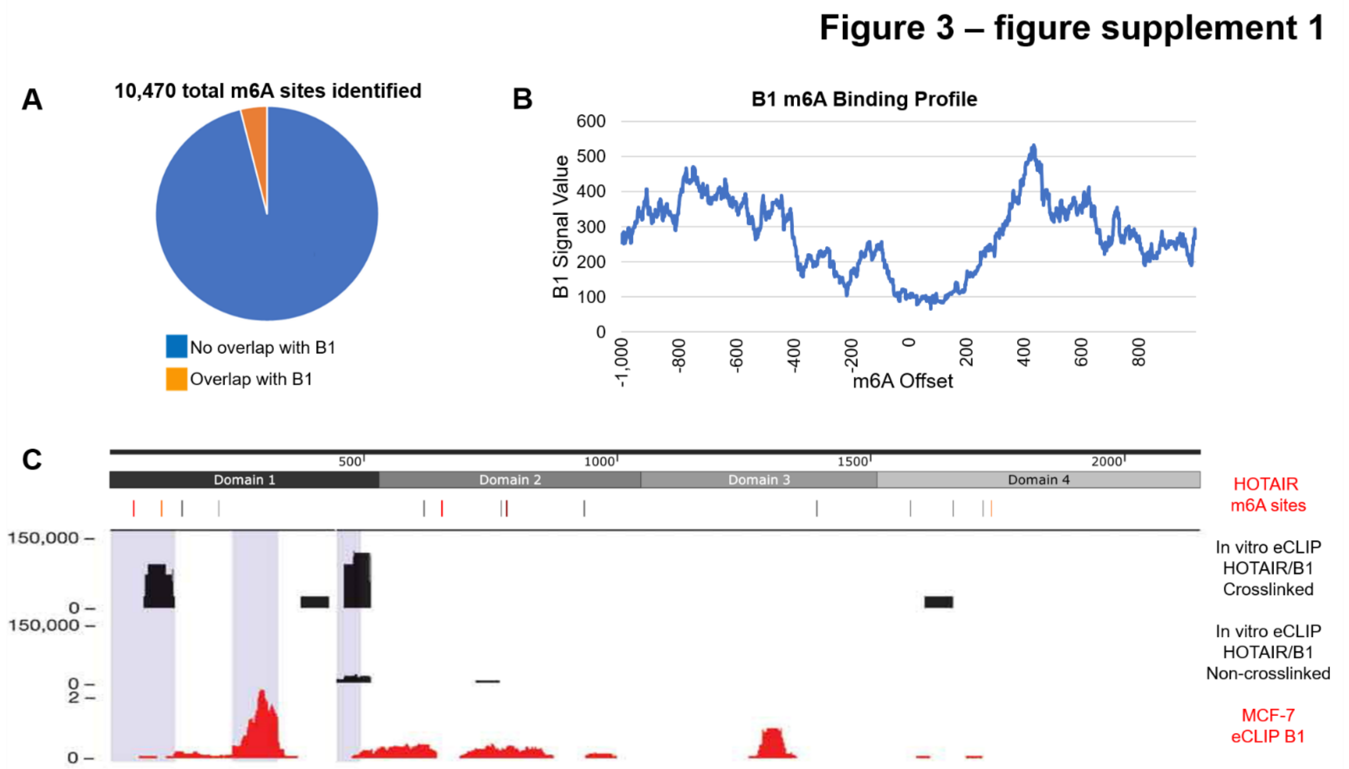


**Figure 3 – figure supplement 1. hnRNP B1 does not directly interact with m6A**. A) Results of a search of 2000 base pair regions surrounding m6A sites for hnRNP B1 eCLIP peaks using set functions. B) hnRNP B1 eCLIP intensity relative to m6A sites that contain overlap with hnRNP B1 binding sites within 1000 nucleotides upstream or downstream. C) Map of HOTAIR m6A sites, *in vitro* eCLIP peaks of hnRNP B1 binding, and hnRNP B1 eCLIP peaks in MCF-7 cells(Nguyen et al., 2018).

#### Figure 4 – figure supplement 1

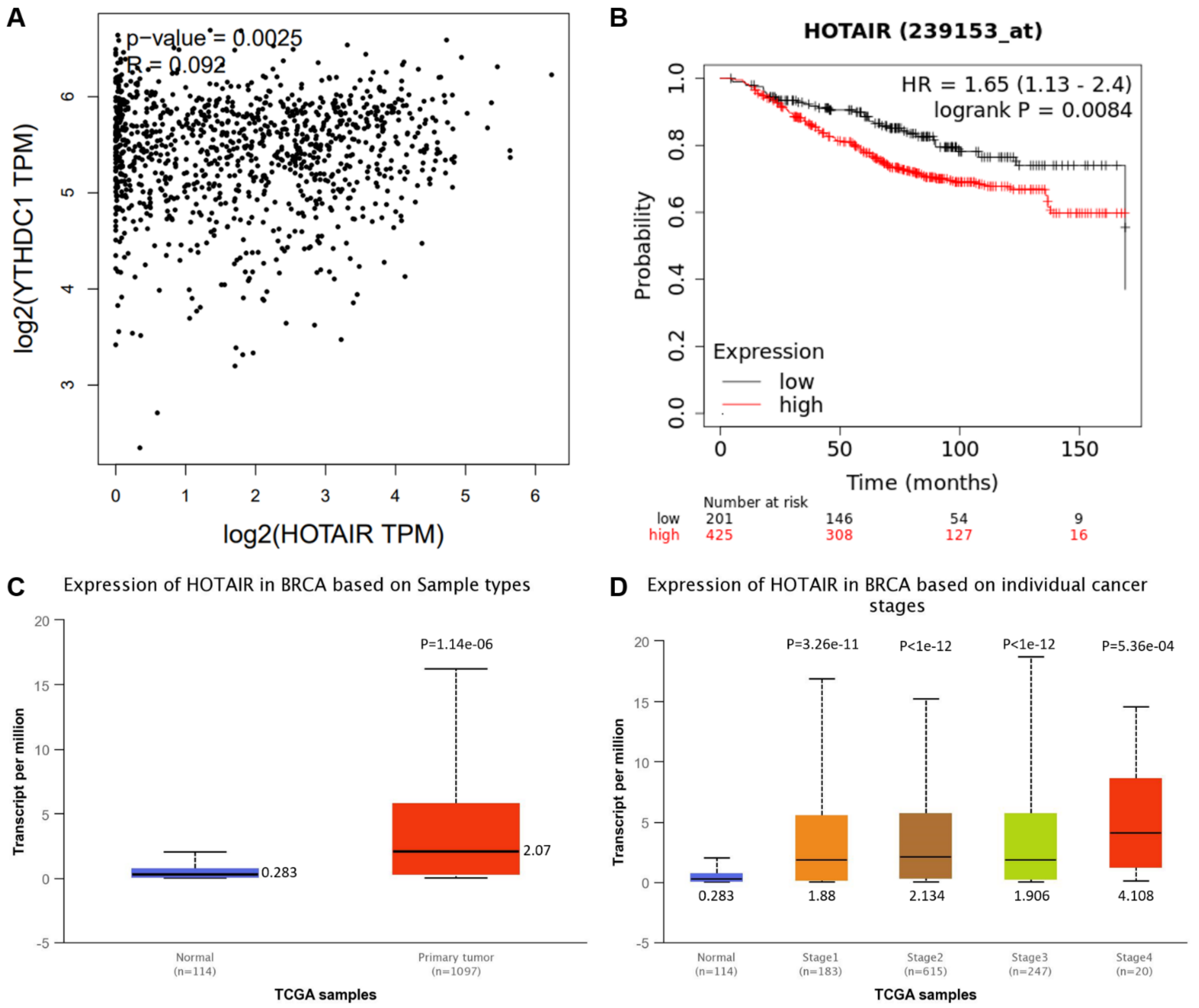


**Figure 4 – figure supplement 1. YTHDC1 and HOTAIR expression correlation in breast cancer. A)** Correlation of *HOTAIR* and *YTHDC1* expression in invasive breast cancer generated from GEPIA2 (Tang et al., 2019). **B)** Kaplan-Meier curves for overall survival of breast cancer patients with high or low expression of *HOTAIR* generated using Kaplan-Meier Plotter(Gyorffy et al., 2010). **C-D)** *HOTAIR* expression in C) normal vs. primary tumor samples and D) normal and stage 1-4 tumor samples. Plots in C-D were generated with UALCAN(Chandrashekar et al., 2017).

**Figure 4 – figure supplement 2**

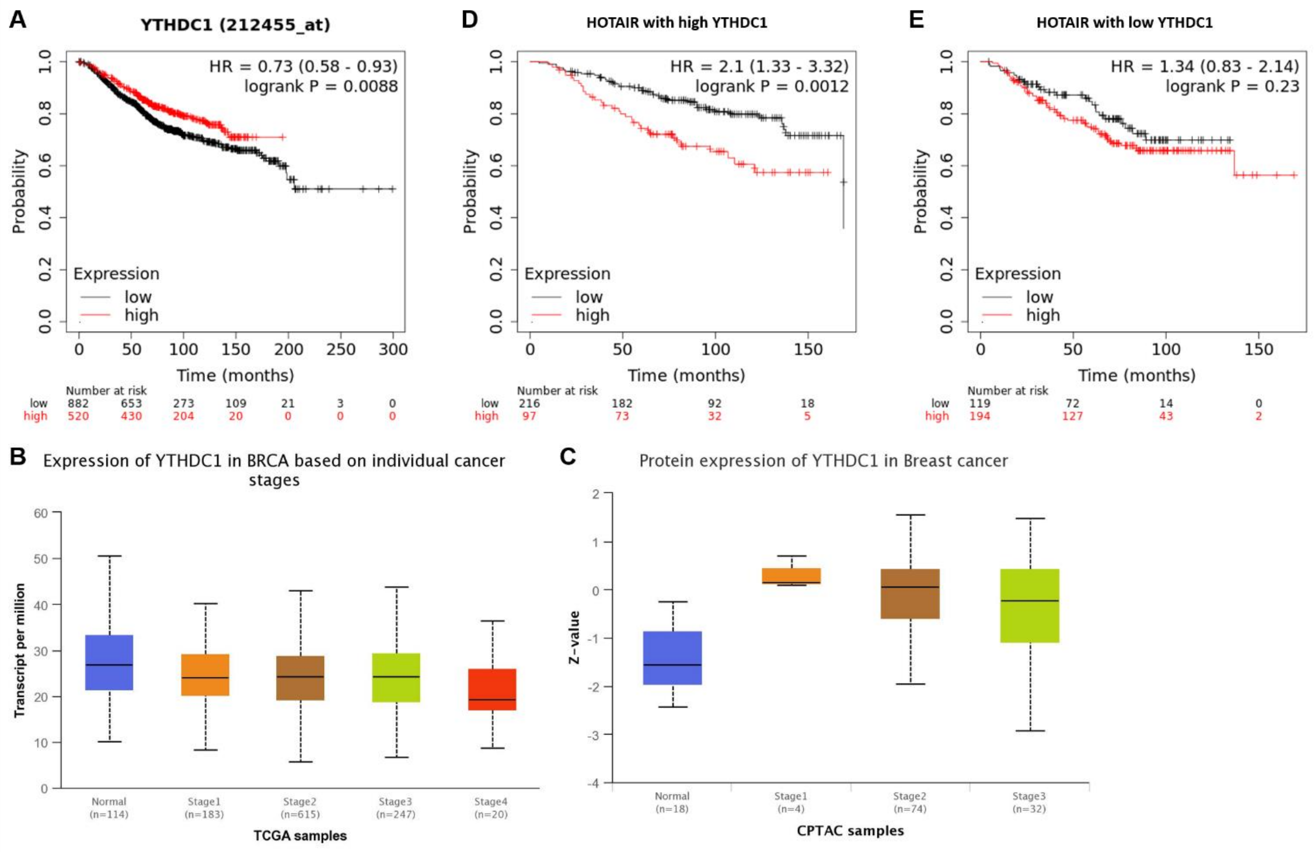


**Figure 4 – figure supplement 2. YTHDC1 expression levels in breast cancer regulate predictive nature of HOTAIR expression. A)** Kaplan-Meier curves for overall survival of breast cancer patients with high or low expression of *YTHDC1* generated using Kaplan-Meier Plotter(Gyorffy et al., 2010). **B-C)** Expression of *YTHDC1* E) RNA and F) protein in normal and stage 1-4 tumor samples. Plots in B-F were generated using UALCAN (Chandrashekar et al., 2017). **D-E)** Overall survival curves for breast cancer patients examining effect of HOTAIR on the background of either D) high or E) low *YTHDC1*.

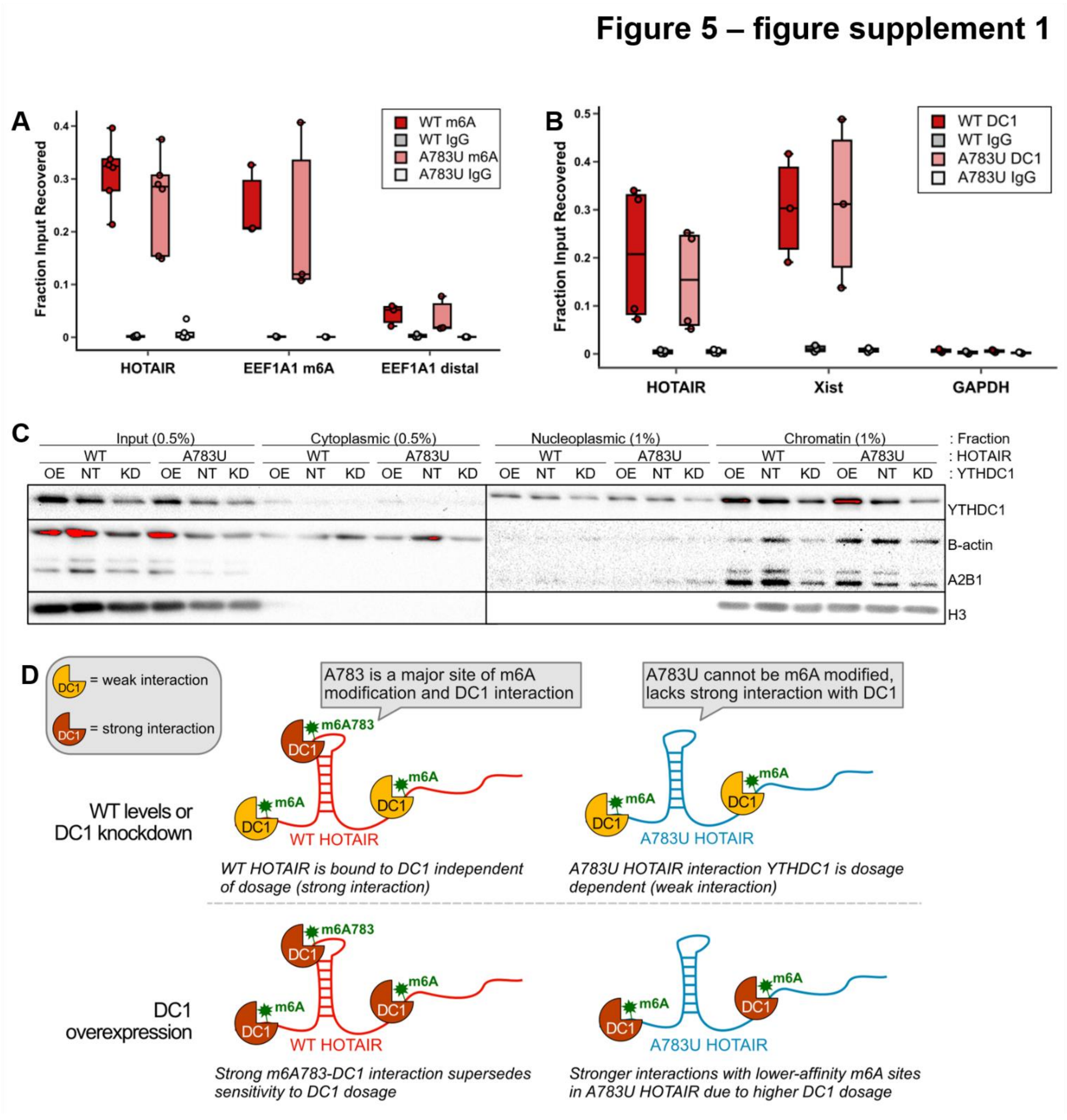


**Figure 5 – figure supplement 1. A783 does not mediate overall m6A modification of overexpressed HOTAIR nor does it regulate interaction with YTHDC1 at other m6A sites *in vivo*, but these m6A sites and YTHDC1 mediates stability of HOTAIR. A)** m6A RIP performed on MDA-MB-231 cells overexpressing WT HOTAIR or A783U HOTAIR. **B)** YTHDC1 RIP performed on MDA-MB-231 cells overexpressing WT HOTAIR or A783U HOTAIR. **C)** Western blot performed on fractionation of MDA-

MB-231 cell lines overexpressing WT or A783U HOTAIR containing overexpression (OE), non-targeting (NT), or knock-down (KD) of YTHDC1. **D) E)** Model for differences observed between WT and A783U HOTAIR upon knockdown and overexpression of YTHDC1. **F)** Schematic of dCasRX-YTHDC1 fusion protein targeted to the HOTAIR transcript.

### Figure 7 – figure supplement 1

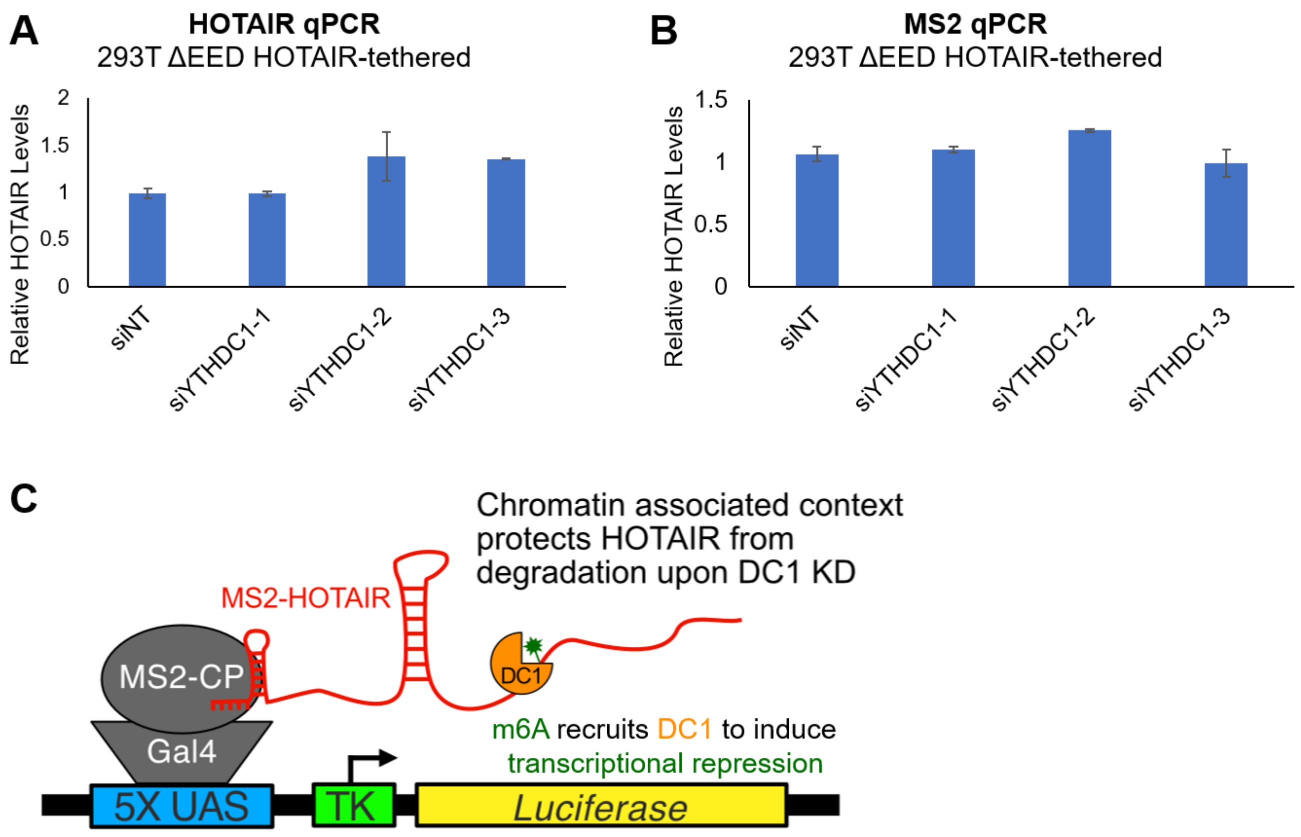


**Figure 7 – figure supplement 1. HOTAIR is stable upon YTHDC1 knockdown in 293T HOTAIR-tethered reporter cell lines. A-B)** qRT-PCR of HOTAIR (A) or MS2 (B) in 293T HOTAIR-tethered cells lacking EED with siRNA knockdown of YTHDC1 or a non-targeting control. **C)** Model for HOTAIR stability in chromatin-tethered context.
