## Supplemental Tables for "A single N6-methyladenosine site in lncRNA HOTAIR regulates its function in breast cancer cells"

### Supplemental File 1: Tables of m6A sites in HOTAIR and ORFs, shRNAs, oligonucleotides, and plasmids used in this study.

| | Experiments | | | | | | | | | | |
|---|---|---|---|---|---|---|---|---|---|---|---|
| HOTAIR m6A site positions | MCF-7 rep1 | MCF-7 rep2 | MCF-7 rep3 | MDA-MB-231 pB-HOTAIR rep1 | MDA-MB-231 pB-HOTAIR rep2 | MDA-MB-231 pB-HOTAIR rep3 | MDA-MB-231 pB-HOTAIR$^{A783U}$ rep1 | MDA-MB-231 pB-HOTAIR$^{A783U}$ rep2 | MDA-MB-231 pB-HOTAIR$^{A783U}$ rep3 | MDA-MB-231 pB-Anti-Luc | 293 HOTAIR-Luc dEED |
| **Nt 48 / 54362413** | | | | **X** | **X** | | **X** | x | **X** | **X** | x |
| **Nt 102 / 54361137** | | | | **X** | x | | x | * | * | | x |
| Nt 143 / 54361096 | | | | * | * | | | | | | x |
| Nt 215 / 54360133 | | | | * | | | x | | | | x |
| Nt 557 / 54357821 | | | | * | | | | x | | | |
| Nt 620 / 54357758 | | | | **X** | x | * | | x | | | * |
| **Nt 655 / 54357723** | | | | **X** | **X** | **X** | **X** | | * | **X** | x |
| **Nt 772 / 54357606** | | | | x | * | x | x | **X** | x | **X** | x |
| **Nt 783 / 54357595** | **X** | * | | **X** | **X** | **X** | | | | **X** | **X** |
| Nt 936 / 54357442 | | | | * | | | x | | | | |
| Nt 1394 / 54356983 | | | | x | | | * | | | | x |
| Nt 1579 / 54356798 | | | | **X** | | | | | | | * |
| Nt 1663 / 54356714 | | | | x | | | | | | * | |
| Nt 1722 / 54356655 | | | | * | | | | * | | | x |
| **Nt 1739 / 54356638** | | | | x | **X** | | **X** | * | | | x |

Legend: **X**=high confidence x=low confidence *=reduced threshold

**Table S1. List of HOTAIR m6A sites by experiment.** Each column represents a single experiment with the cell line noted. Each row is an m6A site detected within HOTAIR and includes Nt # (location within HOTAIR transcript) and chromosome position. Each X represents an m6A site detected in HOTAIR in each experiment. Using thresholding from (Roberts, Porman, & Johnson, 2020), '**X**' represents high confidence m6A sites (≥3 C→T mutations following the m6A site in ≥5% of reads), 'x' represents low confidence sites (≥3 C→T mutations following the m6A site in ≥2.5% of reads), and '*' represents sites called with reduced threshold of at least 2 C→T mutation events detected. HOTAIR m6A site positions in bold were included in the 6x HOTAIR mutant, while the remainder of sites (not including Nt 557) were included in the 14x HOTAIR mutant. Note that MCF-7 replicate 3 was a lower-depth run used as a point of comparison to the higher-depth replicates 1 and 2 in (Roberts, Porman, and Johnson, 2020). HOTAIR read depth in MCF-7 experiments is ~10x less than when overexpressed in MDA-MB-231 cells. m6A site 783 is highlighted in red.

| HOTAIR m6A sites | MCF-7 | MDA-MB-231 pB-HOTAIR | MDA-MB-231 pB-HOTAIR$^{A783U}$ |
|---|---|---|---|
| **Nt 48 / 54362413** | | X | X |
| **Nt 102 / 54361137** | | X | X |
| Nt 143 / 54361096 | | X | |
| Nt 620 / 54357758 | | X | |
| **Nt 655 / 54357723** | | X | X |
| **Nt 772 / 54357606** | | X | X |
| **Nt 783 / 54357595** | X | X | |
| **Nt 1739 / 54356638** | | X | X |

**Table S2. Multiple replicate consensus list of m6A sites in HOTAIR-expressing breast cancer cell lines.** X indicates an m6A site detected in 2+ replicates in the cell line noted.

| Gene / Knockdown or Overexpression | shRNA/ORF Used |
|---|---|
| METTL3 Knockdown | TRCN0000034715, TRCN0000034717 |
| METTL14 Knockdown | TRCN0000015933, TRCN0000015937 |
| WTAP Knockdown | TRCN0000231422, TRCN0000231424 |
| YTHDC1 Knockdown | TRCN0000243987, TRCN0000243989 |
| YTHDC1 Overexpression | ORF clone ccsdBroad304_04559 |

**Table S3. List of shRNAs and ORFs used in this study.**

| Experiment | Forward Oligo | Reverse Oligo |
|---|---|---|
| A783U HOTAIR QuikChange | AG66<br>CGCCCAGAGAtCGCTGGAAAAACCTGAGCGG | AG67<br>CCAGCGaTCTCTGGGCGTTCATGTGGCGAGC |
| A783U HOTAIR pBABE-Puro | AG68<br>CCTAAACCAGCAATTACACCCAAGCTCGTTGGGGCCTAAG | AG69<br>CTGTGCTGGCGAATTCCTACGTACCACCACACTGGGATCCGAAAATGCATCCAGATATTAAT |
| Cloning YTHDC1 into pCDNA3-FLAG | AG64<br>ggtaccGCGGCTGACAGTCGGGAGGAG | AG65<br>gcggccgccCTAATCTTCTATATCGACCTCTC |
| NheI cloning YTHDC1 into pXR002 | AL01<br>gtagctgctagcGACTACAAAGACGATGACGATAAAGGGG | AL02<br>gatgctgctagcTCTTCTATATCGACCTCTCTCCCCTCG |
| HOTAIR gRNA cloning into pXR003 | AL05<br>AAACCCCCGGCACCCGCTCAGGTTTT | AL10<br>AAAAAAAACCTGAGCGGGTGCCGGGG |
| Non-targeting gRNA cloning into pXR003 | AL07<br>AAACCAGAAGCGTACCATACTCACGA | AL 11<br>AAAATCGTGAGTATGGTACGCTTCTG |

**Table S4. Oligonucleotides used for constructing pBABE-Puro-A783U_HOTAIR and dCasRX-YTHDC1.**

| **m6A eCLIP oligonucleotides** | **Sequence** |
|---|---|
| X1a (RNA) | /5Phos/rArUrArUrArGrG rNrNrNrNrN<br><br>rArGrArUrCrGrGrArArGrArGrCrGrUrCrGrUrGrUrArG/3SpC3/ |
| X1b (RNA) | /5Phos/rArArUrArGrCrA rNrNrNrNrN<br><br>rArGrArUrCrGrGrArArGrArGrCrGrUrCrGrUrGrUrArG/3SpC3/ |
| Rand3Tr3 (RNA) | /5phos/rArGrArUrCrGrGrArArGrArGrCrGrUrCrGrUrG/3SpC3/ |
| RiL19 (RNA) | /5phos/rArGrArUrCrGrGrArArGrArGrCrGrUrCrGrUrG/3SpC3/ |
| AR17 (DNA) | /5Phos/NNNNNNNNNNAGATCGGAAGAGCACACGTCTG/3SpC3/ |

**Table S5. Oligonucleotides used for m6A eCLIP**

| Gene | qPCR Forward Primer | qPCR Reverse Primer |
|---|---|---|
| HOTAIR | GGGAACGGGAGTACAGAGAGAATA | GGCACCCGCTCAGGTTT |
| GAPDH | CCGGGAAACTGTGGCGTGATGG | AGGTGGAGGAGTGGGTGTCGCTGTT |
| XIST | AAACCCAACACGAAAAGCAC | GCGGTCACACAGGAAAAGAT |
| EEF1A1 +m6A | CGGTCTCAGAACTGTTTGTTTC | AAACCAAAGTGGTCCACAAA |
| EEF1A1 distal | GGATGGAAAGTCACCCGTAAG | TTGTCAGTTGGACGAGTTGG |
| PTK7 | ACACTTCGTTGCCACATTGAT | CAGCAGGAATACAGCCCAC |
| CDH11 | AGAGGTCCAATGTGGGAACG | GGTTGTCCTTCGAGGATACTGT |
| GRIN2A | TGGCCTCACCGGGTATGATT | CAATGCCGTCCCTCACTCTC |
| SEMA5A | GATCCTGCCATTTACCGAAGC | AGATGACACAAAGTTTGGCTCA |
| SIRPA | ACATGGTCCACCTCAACCG | ACGCTGGCGTACTCTGAGA |
| TP53I11 | GAAGACCCGCAAGATCCTCG | TTTCATTGCCTAAGACCTGGC |
| Luciferase (LucR2) | GCACTGATCATGAACTCCTCTGGATCTAC | GAGAATAGGGTTGGCACCAGCAGCGCAC |

**Table S6. Oligonucleotides used for qPCR.**

<table>
<tr><th>Fragment</th><th>Template</th><th>F Primer</th><th>R Primer</th></tr>
<tr><td>WT HOTAIR D2</td><td>pAJ249</td><td rowspan="2">MB88<br><br>GTAATACGACTCACT ATAGGGAGCCAGAG GAG</td><td rowspan="2">MB89<br><br>CCATATAAACTCCTT AAAGCTTATATTTTA CAGTCC</td></tr>
<tr><td>A783U HOTAIR D2</td><td>pAJ385</td></tr>
<tr><td>RAT-WT D2</td><td>WT HOTAIR D2</td><td rowspan="2">MB22<br><br>TAATACGACTCACTA TAGGG</td><td rowspan="2">MB94<br><br>CGATGGCACGAGTG TAGCTAAACCTCGTG CCGACGTCTAAGGG TTTCCATATAAACTC CTTAAAGCTT</td></tr>
<tr><td>RAT-A783U D2</td><td>A783U HOTAIR D2</td></tr>
</table>

**Table S7. Plasmids and oligonucleotides used for *in vitro* m6A methylation experiments.**
